# A tunnel microtract organ for T cell progenitor homing is formed by neural crest morphogenesis via Sox10-Cdc42 axis

**DOI:** 10.64898/2026.03.05.709688

**Authors:** Fangying Zhao, Jiangyong He, Liqing Xiao, Hang Chen, Zhifan Li, Yafang Lu, Li Duan, Junlong Zhao, Gengrong Chen, Xiaobei Huang, Lingfei Luo, Li Li

## Abstract

Thymic colonization by T cell progenitors (TCPs) is essential for adaptive immunity, yet the guiding tissues remain elusive. Here, we unveiled a tunnel microtract (TMT) as an organ indispensable in TCPs homing in both zebrafish and mouse. Disruption of TMT leads to compromised T cell development. Specifically, the zebrafish TMT were positioned bilaterally beneath the fifth branchial levator muscle, connecting the thymus and kidney. They are semi-coiled, non-vascular, non-lymphatic tubes of epithelial signatures. Impressively, Sox10 activates Cdc42 to promote F-actin remodeling in neural crest cells (NCCs), leading to precise elongation and tight packaging of a low-permeability TMT. Remarkably, a homologous CD34⁺ TMT was observed in mice, which bilaterally enveloped the embryonic thymus and extended into the thyroid cartilage. Sox10-Cdc42 signalling functioned recapitulatively in NCC morphogenesis during its construction. These findings establish TMT as an unappreciated NCC-derived organ in TCPs homing, with implications in T cell development and immune disorders.

## Introduction

The continuous homing of hematopoietic progenitor cells (HPCs) from extrathymic hematopoietic organs to the thymic microenvironment enables lineage commitment, T-cell receptor rearrangement, and selection processes that establish immune competence and self-tolerance^1–3^. This process is initiated at the embryonic stage and is indispensable for establishing and maintaining a functional adaptive immune system. In mice, lymphoid progenitors arise from the yolk sac (YS) on embryonic day 7.5 (E7.5) and the aorta-gonad-mesonephros (AGM) on E9, seeding the fetal liver on E10.5, as a transient reservoir for export to peripheral organs, including the thymus^4^. By E14.5, the bone marrow becomes the major HPC niche, supplanting earlier sources and serving as the primary supply of thymic progenitors in late fetal and postnatal life^5^. Thymic colonization begins on E11.5, with CD45^+^ hematopoietic precursors migrating to the avascular rudiment^6^. This prevascular phase (E11.5–E12.5) proceeds via a blood-independent mechanism: precursors extravasate from adjacent vessels, traverse the perithymic mesenchyme, and cross the epithelial basement membrane, guided by G protein-coupled receptors sensing chemokine gradients (notably CCL21 and CCL25), and regulated by axes such as CXCR4-CXCL12 for mobilization from niches, including the bone marrow^7–9^. Following vascularization later in gestation, progenitors enter the bloodstream predominantly at the corticomedullary junction (CMJ)^10^. Here, circulating HPCs extravasate through specialized P-selectin^+^Ly6C^−^ thymic portal endothelial cells (TPECs) into perivascular spaces before entering the parenchyma, orchestrated by molecular cues including SIRPα-CD47 interactions and TEC-derived chemokines^11–13^. Although post-vascularization homing has been observed in mice, the mechanisms and anatomical structures underlying the initial colonization of the prevascular rudiment remain poorly defined.

Thymus trafficking and T cell development exhibit broad conservation^14–16^. Embryonic studies in zebrafish have revealed the early progenitor origins and homing of hematopoietic cells, based on imaging advances in transparent embryos. By ∼54 hours post-fertilization (hpf), CD41-GFP^low^ progenitors undergo extended mesenchymal migration to the nascent thymus^17^. Time-lapse imaging revealed three distinct extravasation routes for progenitor colonization of the prevascular thymic rudiment^18^. The predominant route featured egress from a trifurcated vascular junction (posterior cardinal vein, primary head sinus, and dorsal aorta; site 1). A secondary pathway involved vessels adjacent to the rudiment (site 2), while a minor rostrocaudal route entailed exit at site 3, followed by rapid parenchymal infiltration. These findings have established a hierarchical map of the vascular-dependent progenitor entry routes that operate prior to thymic vascularization^18^. In the larval stages (beyond 5 days post-fertilization (dpf)), when the kidney marrow becomes the principal hematopoietic organ (analogous to the mammalian bone marrow)^19^, the ontogeny of thymus-settling progenitors, their primary sources, and strategies enabling peripheral-to-thymic homing are largely uncharacterized.

Neural crest cells (NCCs) are transient multipotent cells that arise from the dorsal neural tube during early vertebrate embryogenesis^20^. They undergo epithelial-to-mesenchymal transition (EMT) and migrate to diverse sites, where they differentiate into a broad range of derivatives, including neurons and glia of the peripheral nervous system, melanocytes, craniofacial cartilage and bone, and various mesenchymal lineages such as connective tissues^20, 21^. However, NCCs play conserved yet model-specific roles across species in thymus organogenesis^22^. In mice, NCCs migrate ventrally into the pharyngeal arches around E9.5–E10.5^23^. NCC-derived mesenchymal cells condense around the third pharyngeal pouch-derived thymic primordium to establish an essential pericapsular stromal niche (the perithymic mesenchyme)^24, 25^. Through the secretion of the morphogens FGF7, FGF10, and BMP4, this niche orchestrates TEC proliferation and patterning, facilitates early lymphoid progenitor recruitment via chemokine gradients, and supports vascularization and thymocyte egress^24^. Zebrafish NCCs contribute to thymus morphogenesis by interacting with the pharyngeal endoderm, with Chd7 modulating both the NCC-derived mesenchyme and endoderm-derived TECs^26, 27^. Chd7 deficiency impairs organogenesis and lymphoid progenitor homing, downregulates pharyngeal Foxn1 expression, and causes T-cell lymphopenia (despite increased peripheral hematopoietic stem and progenitor cells)^27^. This T-cell reduction can be partially rescued by Foxn1 overexpression, highlighting the important role of NCCs in establishing a thymic stromal microenvironment in zebrafish.

The high-mobility group (HMG) box transcription factor Sox10, a core member of the SoxE subfamily, along with Sox8 and Sox9, serves as a master regulator of NCC development across vertebrates^28, 29^. It orchestrates cell fate determination, survival, and migration by acting as a pioneer factor in remodeling the chromatin architecture and by directly activating key downstream effectors, such as FoxD3, ErbB3, Ret, and Mitf^30–32^. A critical aspect of this function is the regulation of cytoskeletal dynamics and cellular morphology, which underlie tissue-level patterning^33^. This is achieved through its role in the SOX9/SOX10 signalling axis, which controls the polarized activity of RHOA via the RhoGAP protein DLC1, thereby establishing the front-rear polarity essential for directional migration^34^. Although the roles of Sox10 in the glial and melanocytic lineages are well defined, its potential involvement in driving epithelial-like morphogenesis within neural crest-derived stromal structures remains unclear.

As a Rho family GTPase, Cdc42 functions as a central regulator of F-actin organization and directly controls filament nucleation, branching, and spatial assembly to enable filopodial extension, collective cell migration, and tissue-scale morphogenesis^35–37^. Cycling between the GDP-bound and GTP-bound states, Cdc42 activates N-WASP and the Arp2/3 complex to generate branched actin networks, while engaging PAK kinases to modulate actin turnover and membrane dynamics^37^. Precise spatiotemporal control of Cdc42 by guanine nucleotide exchange factors, GTPase-activating proteins, and guanine nucleotide dissociation inhibitors ensures coordinated actin remodeling, which is essential for the structural integrity of tubular conduits^38^. In epithelial tubulogenesis models, including kidney collecting ducts, lung airways, vascular networks, and neural tube development, Cdc42 deficiency disrupts the F-actin architecture, impairs protrusive activity, blocks lumen initiation, and results in collapsed or cystic tubular structures^39^.

In the present study, we have identified the tunnel microtract (TMT) as a previously unrecognized NCC-derived organ that serves as a dedicated conduit for embryonic T cell progenitor (TCP) homing in both zebrafish and mice. In zebrafish, the TMT (zTMT) consists of a tubular monolayer of *fli1*^+^*flk1*^-^*lyve*^-^ cells situated beneath the fifth branchial levator muscle connecting the thymus and kidney marrow. These cells undergo a morphological transition from rounded to spindle-shaped of *sox10*⁺ NCCs, establishing a low-permeability barrier that ensures unidirectional progenitor trafficking and priming. Single-cell transcriptomics and lineage tracing revealed their NCC origin and stepwise differentiation into a defined *hoxd11a*⁺*tbx5a*⁺ guidance epithelium. In mice, a homologous CD34⁺ NCC-derived TMT (mTMT) surrounds the embryonic thymus and extends to the thyroid cartilage. Flattened CD34⁺ cells, which are distinct from classical vascular endothelial or thymic epithelial cells, form a tubular conduit enclosing lymphocytes and recapitulating the zTMT architecture. Impressively, Sox10-dependent Cdc42 activation in F-actin cytoskeletal remodeling is conservatively essential for NCC morphogenesis in both zTMT and mTMT formation, supporting the thymic homing of TCPs. These findings therefore establish the TMT as a previously unappreciated definitive NCC-derived organ that is indispensable in the embryonic thymic homing of TCPs and provides cues for understanding adaptive immunity and related disorders.

## Results

### A tunnel microtract directs T cell progenitor homing to the thymus in zebrafish embryos

T cell progenitors (TCPs) originate from extrathymic hematopoietic organs and migrate to the thymus for maturation^10^; however, the organs mediating this homing process remain poorly defined. To this end, we labelled *coro1a*^+^ cells harboring the *ikzf1*^+^ TCP signature (Extended Data Fig. 1a) within the aorta-gonad-mesonephros (AGM), caudal hematopoietic tissue (CHT), and kidney (a functional homologue of bone marrow in mammals) of *Tg(coro1a:Kaede)* larvae by photoconversion (Fig. 1a and Extended Data Fig. 1b)^40–42^. Time-lapse imaging revealed that approximately five red *coro1a*-Kaede^+^ cells from the kidney, but not AGM or CHT, primarily migrated to the thymus through a defined path beneath the fifth branchial levator (BL5) muscle^43^ during the 300-minute period (Fig. 1a-c, Extended Data Fig. 1b and Supplementary Video 1). This migratory pattern was intensified by a chain of coro1a-DenNTR^+^ cells during T-cell recovery in metronidazole (MTZ) treated *Tg(coro1a:DenNTR)* larvae (Fig. 1d)^40^. Structure analysis of the migration path of *coro1a^+^*cells by hematoxylin and eosin (HE) staining and electron microscopy revealed a linear duct-like structure composed of single-cell-layer flattened cells of limited motile cilia with a ∼5 μm lumen, which contained lymphoid progenitor-like cells (Fig. 1e-g). Examination of the reporter lines, including *Tg(fli1a:eGFP)* for endothelial/epithelial cells, *Tg(flk1:mCherry)* for blood vessels, *Tg(prox1:RFP)* for lymphatic vessels, and *Tg(cdh17:DsRed)* for renal tubules, revealed that this path could be labelled by fli1a-eGFP, but not by flk1-mCherry, prox1-RFP, or cdh17-DsRed (Fig. 1h,i). We further excluded cartilage (*Tg(snorc:GFP)*, II-II6B3 antibody) and muscle identity (S58, F310 antibodies) from this pathway (Fig. 1j-l). The fli1a-eGFP^+^ cells precisely outlined the conduit through which the coro1a-DsRed^+^ TCPs actively migrated toward the thymus (Fig. 1m, Extended Data Fig. 1c and Supplementary Video 2). However, limited macrophages (mpeg1-DsRed^+^)/granulocytes (lyz-DsRed^+^), and no erythrocytes (gata1a-DsRed^+^) were observed in the flila-eGFP^+^ structure (Fig. 1n and Extended Data Fig. 1d). These findings reveal a previously uncharacterized luminal tunnel microtract (TMT) defined by *fli1a*⁺*flk1*⁻*prox1*⁻ expression that mediates the thymic homing of TCPs from the kidney (Fig. 1o).

**Fig. 1.**
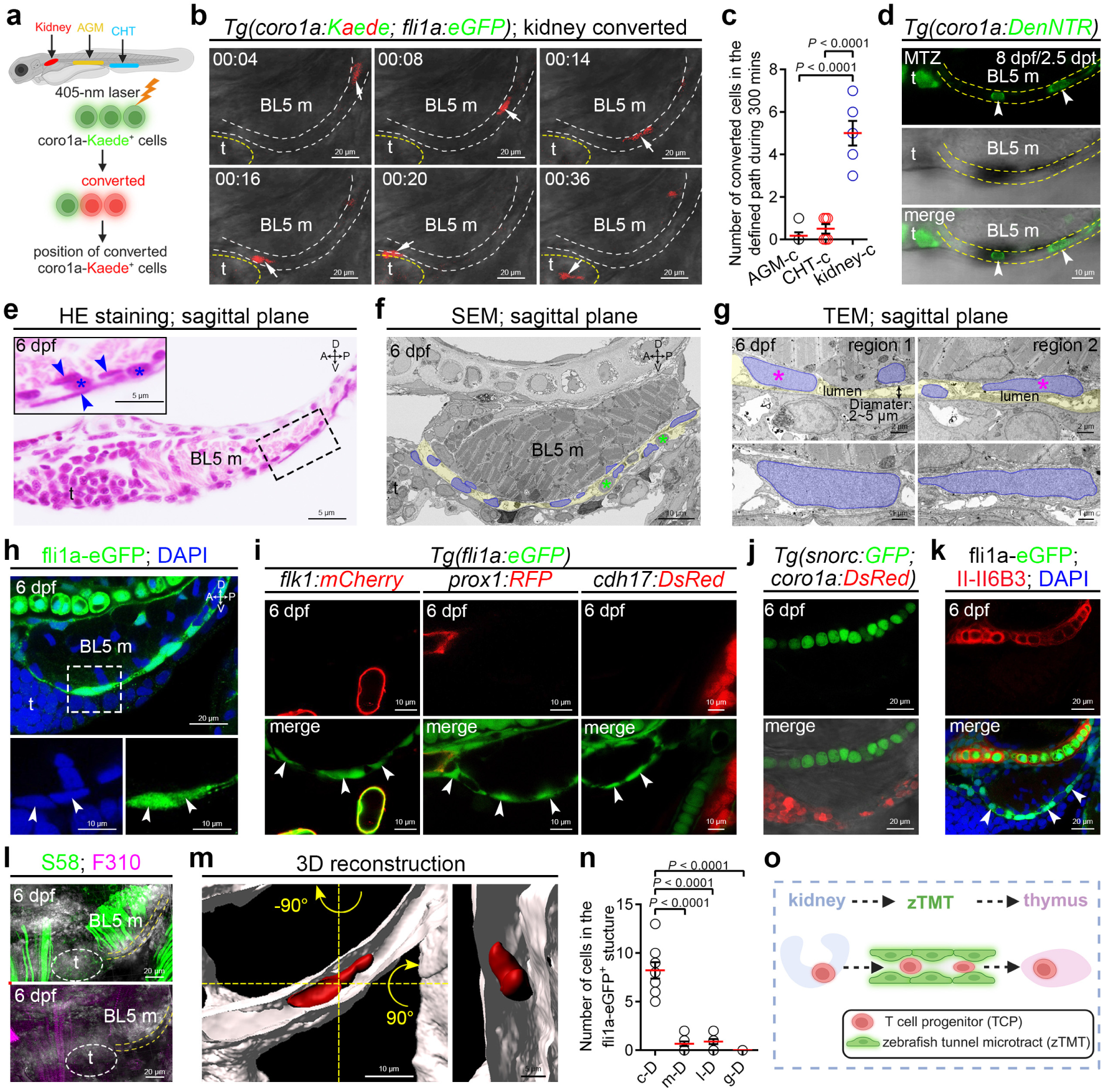
Identification of a *fli1a^+^*zebrafish tunnel microtract (zTMT) mediating thymic homing of T cell progenitors (TCPs). **a**, Schematic diagram of photoconversion of coro1a-Kaede^+^ cells in zebrafish larvae at 5 dpf. **b**, Time-lapse images showing photoconverted (red) coro1a-Kaede^+^ cells (white arrows) migrating from the kidney to the thymus (outlined by yellow dashed lines) along a defined path (outlined by white dashed line). **c**, Quantification of photoconverted coro1a-Kaede^+^ cells migrating from AGM, CHT, and kidney to the thymus along the path over 300 min of imaging. **d**, Confocal images of coro1a-DenNTR^+^ cells (white arrowheads) aligned along the defined path (outlined by yellow dashed lines) in 8 dpf/2.5 dpt MTZ-treated zebrafish larvae. **e**, Representative HE staining of sagittal sections of the path structure. Enlarged view of the black dotted box is shown in the upper left panel insert. **f-g**, SEM (**f**) and TEM (**g**) images of sagittal sections showing the path structure. Cell nuclei are pseudo-colored in blue; the lumen is highlighted in yellow. Enlarged views of magenta asterisks in **g** are shown in the bottom panels. **h**, Confocal images showing fli1a-eGFP^+^ cells in the define path region at 6 dpf. Enlarged views of the white dotted box are shown in the bottom panels. **i**, Confocal images showing the absence of flk1-mCherry^+^ blood vessels (left), prox1-RFP^+^ lymphatic vessels (middle), and cdh17-DsRed^+^ nephridium (right) in the fli1a-eGFP^+^ path region. **j-k**, Confocal images showing the absence of snorc-GFP^+^ (**j**) and II-II6B3^+^ (**k**) cartilage in the path region at 6 dpf. **l**, Confocal images showing the absence of S58^+^ slow contracting muscle (green) and F310^+^ fast myosin (magenta) in the path region (yellow dashed lines) at 6 dpf. **m**, 3D reconstruction showing a coro1a-DsRed^+^ cell (red) with elongated morphology migrating inside the tubular fli1a-eGFP^+^ structure (white). Orthogonal view is shown at right. **n**, Quantification of the number of coro1a-DsRed^+^ (c-D^+^), mpeg1-DsRed^+^ (m-D^+^), lyz-DsRed^+^ (l-D^+^), and gata1a-DsRed^+^ (g-D^+^) cells within the fli1a-eGFP^+^ path at 6 dpf. **o**, Schematic diagram of a zTMT guiding TCPs homing from the kidney to the thymus, created with BioRender.com. t, thymus; BL5 m, the fifth branchial levator muscled; dpf, day post-fertilization; dpt, day post-treatment; MTZ, metronidazole; HE, hematoxylin and eosin; A, anterior; P, posterior; D, dorsal; V, ventral. Blue arrowheads and asterisk in **e** mark path-constituting cells and cells inside the path, respectively. Green asterisks in **f** mark cells located within the path. White arrowheads in **h**,**i**,**k** mark path-constituting fli1a-eGFP^+^ cells. Thymus in **l** is outlined by white elliptical dashed lines. Each dot in **c**,**n** represents an individual zebrafish larva. Error bars represent mean ± SEM. Unpaired two-tailed Student’s t-test. *P* values are included in the graphs.

### The configuration of the zebrafish tunnel microtract (zTMT) reveals an adaptive lamellar enclosed architecture

The zTMT structure was investigated. fli1a-eGFP^+^ cells formed an arc-shaped arrangement, with the rostral (anterior) end extending dorsally toward the cranial tip of the thymus and the caudal (posterior) end curving back along the tail of the BL5 muscle (Extended Data Fig. 2a-c). Transverse sectioning was performed at four key anatomical landmarks along the rostrocaudal axis: the rostral end of the thymus (site 1), thymic mid-region (site 2), muscular mid-region (site 3), and caudal end of the muscle (site 4). The fli1a-eGFP⁺ cells displayed bilateral symmetry across the transversal plane. At sites 1, 2, and 3, an arcuate morphology closely enveloping the dorsal thymic surface was observed, with maximal coverage observed at site 2 (Extended Data Fig. 2d). In contrast, at sites 3 and 4, the fli1a-eGFP⁺ cells adhered tightly to the outer layers of the BL5 muscle, showing the broadest interface at site 3 (Extended Data Fig. 2d). Quantitative assessment of fli1a-eGFP⁺ signals (indicated by white dashed lines) revealed that the average cross-sectional lengths of the fli1a-eGFP⁺ zTMT were approximately 48.1 μm, 77.8 μm, 110.2 μm, and 70.7 μm at sites 1–4, respectively (Extended Data Fig. 2e). Sagittal sections along the rostral–caudal axis revealed that fli1a-eGFP⁺ cells form arc-shaped contours across multiple *z*-axis depths (15 μm, 30 μm, 45 μm, and 60 μm), closely conforming to the underlying geometry of adjacent structures, including the BL5 muscle, thymus, and pharyngeal arches (Extended Data Fig. 2f). Quantitative analysis of the fli1a-eGFP⁺ signal expanse (demarcated by white dashed lines) from the caudoventral margin of the BL5 muscle toward the dorsorostral thymic region (at *z* = 15, 30, and 45 μm) or the pharyngeal arch (at *z* = 60 μm) yielded average sagittal lengths of 163.6 μm, 223.6 μm, 125.0 μm, and 93.9 μm, respectively (Extended Data Fig. 2g). Notably, coro1a-DsRed⁺ cells were consistently detected across all four *z*-planes, suggesting that this flattened, tunnel-like architecture provides multiple parallel “lanes” that facilitate efficient thymic colonization by TCPs (Extended Data Fig. 2f). Integrating these cross-sectional and sagittal observations, we reconstructed a 3D lateral view (50 μm depth of field) using the *Tg(fli1a:eGFP; coro1a:DsRed)*, incorporating horizontal (−180°) and vertical (+180°) rotations to fully resolve spatial topology, which discloses that the zTMT adopts a flattened, lamellar morphology arranged in an irregular arc shape (Extended Data Fig. 2h,I and Supplementary Video 3). To determine whether this structure constituted an enclosed lumen, we performed Alexa647-Dextran microinjections (Extended Data Fig. 2j). When the dye was delivered into tissues between the posterior end of the BL5 muscle and the region beneath the otic vesicle (ov), two parallel dextran streams flanking a layer of fli1a-eGFP^+^ cells appeared at 0.5 hours post-injection (hpi) (Extended Data Fig. 2k), indicating the presence of a tubular wall. In contrast, direct injection of Dextran into the lumen resulted in dye retention within the central cavity at 0.5 hpi (Extended Data Fig. 2j,l), suggesting this structure possesses a genuine luminal architecture. Together, these observations indicate that the *fli1a*^+^*flk*^-^*prox1*^-^ luminal duct adopts a semi-circular, lamellar conformation composed of a monolayer of cells, which are irregularly arranged to interface with the surrounding muscles, thymus, and proximal tissues.

### TCPs predominantly enter the zTMT through its caudal terminus

Intensive anatomical analysis revealed that fli1a-eGFP^+^ zTMT extended rostrally over the dorsal surface of the thymus, guiding coro1a-DsRed^+^ TCPs within its lumen toward thymic colonization, while maintaining spatial separation from cells in the thymic parenchyma (Extended Data Fig. 2b,f). At the caudal end of the zTMT, the fli1a-eGFP^+^ signals delineated an elliptical compartment at the posterior terminus of the BL5 muscle. This region was referred to as the caudal end compartment (CEC) (Extended Data Fig. 2c). The CEC receives photoconverted coro1a-Kaede^+^ cells exiting the kidney marrow prior to their entry into the zTMT, as confirmed by *z*-axis imaging and rotated 3D reconstruction (Extended Data Fig. 3a,b and Supplementary Videos 4 and 5). Consistent with these observations, lateral-view HE staining and imaging indicated that the caudal end of the zTMT forms a pouch-like structure composed of a single cell layer situated close to the kidney, where several cells are sparsely distributed in the mesenchymal space for *coro1a*^+^ cell entry (Extended Data Fig. 3c). We performed time-lapse imaging of targeted coro1a-Kaede⁺ cells surrounding the CEC of the zTMT in 4 days post-fertilization (dpf) *Tg(fli1a:eGFP; coro1a:Kaede)* embryos in order to map TCPs entry routes with high spatial resolution (Extended Data Fig. 3d). A total of 56 red-labelled TCPs from 18 embryos were tracked over a 300-minute window. The majority (69.64%; 39/56) entered the zTMT through the CEC and migrated toward the thymus within the lumen (Extended Data Fig. 3e,h and Supplementary Video 6). This was referred to as the CEC route (CR). However, a smaller fraction (10.71%; 6/56) accessed the zTMT via muscle-associated entry site route (MESR) (Extended Data Fig. 3f,h and Supplementary Video 7). These sites correspond to discrete openings along the segment where zTMT intimately contacts the BL5 muscle (Extended Data Fig. 3f). The remaining cells (19.64%; 11/56) did not enter the lumen, but instead migrated along the ventral exterior route (VER) of the zTMT (Extended Data Fig. 3g,h and Supplementary Video 8). These results therefore indicate that TCPs migrate to the thymus via the zTMT, highlighting the zTMT as a dedicated anatomical conduit that supports the thymic homing of TCPs. The CR is the major route through which TCPs enter the zTMT lumen via the CEC entry site (Extended Data Fig. 3i).

### zTMT functions as a pre-thymic niche for early T-lineage priming

Time-lapse imaging of *Tg(coro1a:Kaede; rag2:DsRed)* embryos revealed that coro1a-Kaede⁺ thymic colonization precursors migrating within the zTMT lumen progressively upregulated rag2-DsRed as they approached the thymus (Fig. 2a,b and Supplementary Video 9). To resolve the spatial dynamics of T-lineage priming along this migratory route, we occluded the CEC in 4 dpf embryos using an ammonolysis-based tetra-PEG hydrogel^44^ sealant with rapid gelation and tissue adhesion (Fig. 2c). For visualization, Cy5-NH_2_ was conjugated to PEG-SG, and sequential injections of PEG-SG/Cy5-NH_2_ followed by PEG-NH_2_ were targeted to the CEC (Fig. 2c). Cy5 fluorescence indicated successful lumen sealing, leading to substantial coro1a-DsRed^+^ cell accumulation caudal to the obstruction two days later, with markedly reduced coro1a-DsRed^+^ signals in the thymus compared to controls (Fig. 2d,e,g). The thymic rag2-DsRed^+^ and *rag1^+^*T cell populations were significantly diminished after occlusion at 6 dpf (Fig. 2f,g). Using this assay, we profiled the cell signatures by selectively labelling and collecting the coro1a-Kaede⁺ cells in three distinct compartments after hydrogel-assisted photoconversion (Fig. 2h). These included the thymus, zTMT lumen (hereafter referred to as intra-zTMT), and the periphery of the zTMT (defined as extra-zTMT). Approximately 500 red-converted cells from each compartment were isolated by fluorescence-activated cell sorting (FACS) for parallel RNA sequencing (Fig. 2h). Differentially expressed gene (DEGs) analysis revealed that genes associated with T cell development showed a stepwise increase in expression along the migratory trajectory from extra-zTMT to intra-zTMT, and finally to the thymus (Fig. 2i). Notably, the transcriptional profile of intra-zTMT cells was markedly more similar to that of thymocytes than that of extra-zTMT cells (Fig. 2i). Although transcript levels of mature T cell-associated genes, including *rag1*, *rag2*, *smarca5*, and *ccs*^45–47^, remained lower than those in the thymus, they were dramatically elevated relative to those in extra-zTMT cells (Fig. 2i). Similarly, key early T lineage markers, including *cd81b*, *esr2b*, *foxp3a*, and *adat1*^48–51^, were significantly upregulated in intra-zTMT cells compared to in extra-zTMT cells, closely resembling their expression patterns in the thymus (Fig. 2i). Gene Ontology (GO) enrichment analysis of genes upregulated in intra-zTMT cells relative to extra-zTMT cells highlighted biological processes, including T cell differentiation in the thymus (GO:0033077), regulation of T cell proliferation (GO:0042129), and adaptive immune responses (GO:0002250) (Fig. 2j). Collectively, these findings indicate that the zTMT is a pre-thymic niche that promotes early T-lineage priming during TCP migration.

**Fig. 2.**
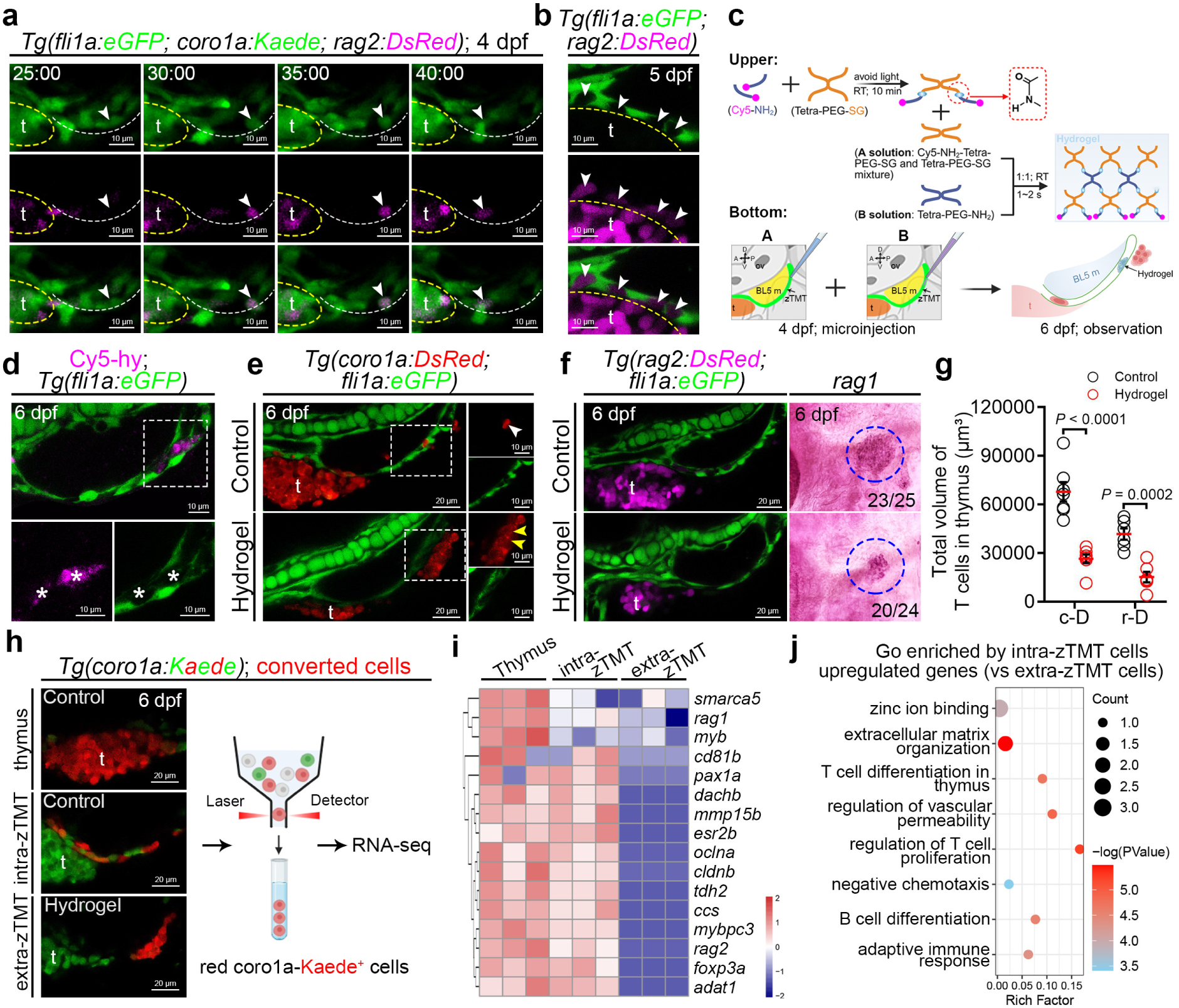
zTMT is essential for early T-lineage priming during TCPs migration. **a,** Time-lapse images showing TCPs gradually expressing rag2-DsRed signals (white arrowheads) within the zTMT (outlined by white dashed line) lumen at 4 dpf. **b**, Confocal images showing rag2-DsRed^+^ cells (white arrowheads) within the fli1a-eGFP^+^ zTMT lumen at 5 dpf. **c**, Schematic diagram illustrating occlusion of zTMT using a 3D hydrogel formed by covalent crosslinking between tetra-PEG-NH_2_ and tetra-PEG-SG. **d**, Confocal images showing Cy5-labelled hydrogel (white asterisks) within the fli1a-eGFP^+^ zTMT lumen at 6 dpf. Enlarged views of the white dashed box is shown in the bottom panels. **e**, Confocal images showing accumulation of coro1a-DsRed^+^ cells (yellow arrowheads) outside the caudal region of zTMT in Cy5-hydrogel-treated larvae, compared with smooth migration of a coro1a-DsRed^+^ cell (white arrowhead) in controls. Enlarged views of boxed regions are shown in the right panels. **f**, Confocal images showing rag2-DsRed^+^ cells within the thymus at 6 dpf following control and hydrogel treatment (left panels). Whole-mount *in situ* hybridization (WISH) analysis of *rag1* in the thymus (blue dashed circles) of control and hydrogel-treated zebrafish at 5 dpf (right panels). **g**, Quantification of coro1a-DsRed^+^ (c-D^+^) and rag2-DsRed^+^ (r-D^+^) signal volumes in the thymus of control and hydrogel-treated zebrafish at 6 dpf. **h**, Experimental workflow for RNA-seq. Left panels show photoconverted red coro1a-Kaede^+^ cells in the thymus, intra-zTMT and extra-zTMT. **i**, Heatmap showing differential expression of mature T cells-associated genes in photoconverted red coro1a-Kaede^+^ cells from thymus, intra-zTMT and extra-zTMT. Color bar represents normalized gene expression levels. **j**, Gene Ontology (GO) analysis of biological processes enriched among up-regulated genes in intra-zTMT cells. Enrichment *P* values were calculated with Fisher’s exact test with FDR correction for multiple testing. Thymus in **a** is outlined by yellow dashed lines. Each dot in **g** represents an individual zebrafish larva. Error bars represent mean ± SEM. Unpaired two-tailed Student’s t-test. *P* values are included in the graphs. Schematic diagrams in **c**,**h** were created with BioRender.com.

### zTMT cell represents a specialized *hoxd11a*^+^*tbx5a*^+^ guidance epithelium

Given the critical role of zTMT as a homing organ for TCPs in T cell lymphopoiesis, we investigated its molecular signature. To this end, we sorted fli1a-eGFP^+^flk1-mCherry^−^ cells from the zTMT niche (peri-ocular region to yolk sac) of *Tg(fli1a:eGFP;flk1:mCherry)* zebrafish at 2, 4, 6, and 8 dpf, respectively, for single-cell RNA sequencing (scRNA-seq) (Extended Data Fig. 4a). Following quality control and UMAP clustering, 13,339, 16,754, 6,521, and 26,316 fli1a-eGFP^+^ cells were resolved into 14 clusters and annotated using lineage-specific markers (Extended Data Fig. 4b-d). Three representative genes from each cluster were selected for whole-mount in situ hybridization (WISH) and fluorescence in situ hybridization (FISH) examination (Extended Data Fig. 4e). The results revealed that factors in cluster 12, including *clic2*, *hoxd11a*, and *tbx5a*, precisely recapitulated the zTMT pattern and colocalized with fli1a-eGFP^+^ cells (Fig. 3a and Extended Data Fig. 4d-f). Furthermore, we observed the zTMT patterns of *hoxd11a,* a homeodomain transcription factor of the Hox gene family with segmental identity along the anterior-posterior axis, and *tbx5a*, a conserved T-box transcription factor essential for forelimb/pectoral fin specification and heart chamber formation^52, 53^. Accordingly, CRISPR/Cas9-mediated knock-in of a Gal4-UAS-P2A-nfsB-mCherry cassette into the endogenous *hoxd11a* locus and generation of a *tbx5a* promoter (6.3 kb)-driven mCherry transgenic line yielded viable, fertile zebrafish (Extended Data Fig. 4g-i). Both hoxd11a-nfsB-mCherry^+^ and tbx5a-mCherry^+^ cells colocalized with fli1a-eGFP^+^ zTMT structures (Fig. 3b,c). Transcriptomic analysis of cluster 12 revealed significant enrichment for morphogenesis of an epithelium (GO:0002009) and epithelial structure maintenance (GO:0010669), with high expression of canonical epithelial markers (*epcam*, *krt18a.1*)^54, 55^, as validated by WISH and FISH (Fig. 3d,e). Notably, zTMT cells lacked expression of the thymic epithelial marker *foxn1*^56^ (Extended Data Fig. 4j,k). These results define zTMT as a specialized epithelial structure with a unique molecular signature.

**Fig. 3.**
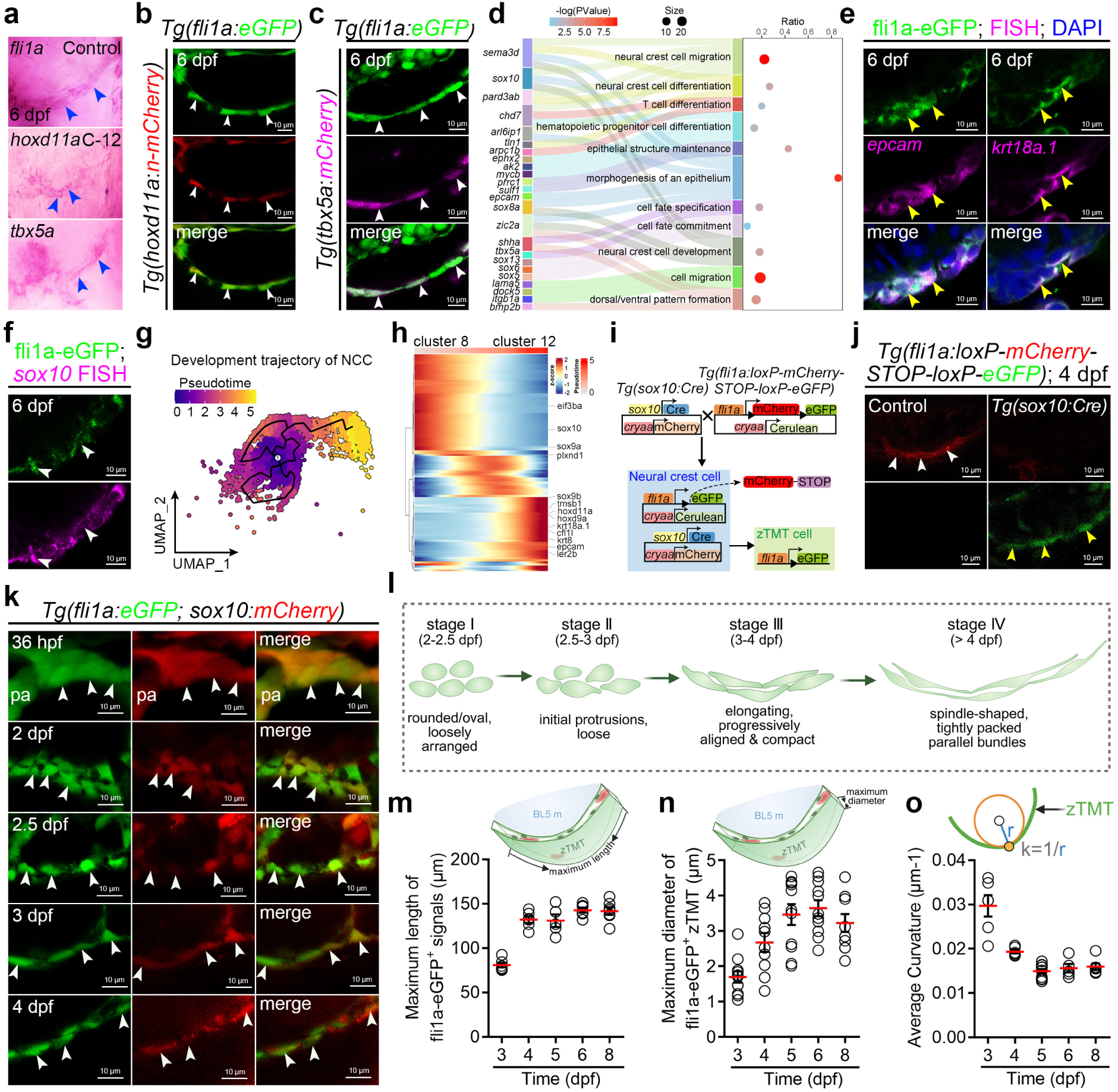
zTMT consists of epithelial cells originated form neural crest cells (NCCs). **a**, WISH signals (blue arrowheads) of *fli1a* and cluster 12-enriched genes (*hoxd11a* and *tbx5a*) in the zTMT region at 6 dpf. **b-c**, Confocal images showing co-expression of hoxd11a-n-mCherry^+^ signals (red, **b**) and tbx5a-mCherry^+^ signals (magenta, **c**) with fli1a-eGFP^+^ zTMT cells at 6 dpf. **d**, GO analysis of biological processes enriched among up-regulated genes in cluster 12. **e**, Confocal images of fluorescence *in situ* hybridization (FISH) for *epcam* (left) and *krt18a.1* (right) mRNA showing colocalization with fil1a-eGFP^+^ zTMT cells at 6 dpf. **f**, Confocal images of FISH for *sox10* mRNA (magenta) showing colocalization with fli1a-eGFP^+^ zTMT cells (green) at 6 dpf. **g**, Pseudotime trajectory inference of NCCs showing developmental routes from cluster 8 to cluster 12. **h**, Heatmap showing the dynamic expression patterns of genes associated with NCC development and epithelial markers along the pseudotime. **i**, Schematic diagram of the lineage tracing strategy using specific transgenic lines to demonstrate the *sox10*^+^ NCC origin of *fli1a*^+^ zTMT cells. **j**, Confocal images showing expression of sox10-mCherry (white arrowheads) and fli1a-eGFP (yellow arrowheads) in 4 dpf *Tg(fli1a:loxP-mCherry-loxP-eGFP)* and *Tg(sox10:Cre;fli1a:loxP-mCherry-loxP-eGFP)* zebrafish. **k**, Confocal images depicting the pattern of sox10-mCherry^+^ signals in the fli1a-eGFP^+^ zTMT region at progressive developmental stages: 36 hpf, 2 dpf, 2.5 dpf, 3 dpf and 4 dpf. **l**, Schematic illustration of the stereotyped morphogenetic program of zTMT. **m-o**, Quantification of the maximum length (**m**), maximum diameter (**n**), and average curvature (**o**) of the fli1a-eGFP^+^ zTMT from 3 dpf to 8 dpf. hpf, hour post-fertilization. White arrowheads in **b**,**c**,**f**,**j**,**k** and yellow arrowheads in **e** indicate overlapping signals. Each dot in **m-o** represents an individual zebrafish larva. Error bars represent mean ± SEM. Unpaired two-tailed Student’s t-test. *P* values are included in the graphs. Schematic diagrams in **i**,**l**,**m-o** were created with BioRender.com

### Neural crest cells give rise to the zTMT through a defined morphogenetic program

Next, we investigated the developmental origin of the specialized epithelial cells that constitute the zTMT. GO enrichment analysis of cluster 12 indicated significant associations between neural crest cell differentiation (GO:0014033), neural crest cell migration (GO:0001755), and neural crest cell development (GO:0014032) (Fig. 3d). Consistent with this, neural crest–associated transcripts, including *sox10* and *sox9a*^29^, were robustly expressed in zTMT cells in cluster 12 (Extended Data Figs. 4d and 5a). The WISH and FISH, coupled with immunofluorescence staining, indicated precise colocalization of *sox10* mRNA with fli1a-eGFP⁺ zTMT cells (Fig. 3f and Extended Data Fig. 5b). Beyond cluster 12 cells, cluster 8 cells similarly expressed *sox10* and *sox9a*, corroborating a shared neural crest cell (NCC) identity for both populations (Extended Data Figs. 4d and 5a), leading us to investigate the relationship between these two clusters through trajectory inference. Pseudotime trajectory inference with Monocle3 indicated an organized, branched differentiation route from the root state (cluster 8) to the terminal state (cluster 12) (Fig. 3g). In addition, genes related to NCC development, such as *sox10*, *sox9a,* and *eif3ba*^57^, exhibited reduced expression patterns along the trajectory from cluster 12 to cluster 8, whereas epithelial signature genes (including *epcam*, *krt18a.1*, and *cfl1l*) showed a remarkable enhancement in expression (Fig. 3h). Furthermore, we employed Cre/loxP-based lineage tracing using *Tg(sox10:Cre; cryaa:mCherry)* and *Tg(fli1a:loxP-mCherry-STOP-loxP-eGFP; cryaa:Cerulean)* fish (Fig. 3i and Extended Data Fig. 5c). Confocal imaging at 4 dpf indicated eGFP labelling of zTMT cells derived from *sox10⁺*cells (Fig. 3j). To this end, we generated *Tg(sox10:NTR-mCherry)* zebrafish and ablated *sox10⁺* cells with MTZ from 3 dpf (Extended Data Fig. 5d). This treatment markedly reduced sox10-NTR-mCherry⁺ and fli1a-eGFP⁺ zTMT cells, disrupted epithelial organization, and severely diminished thymic coro1a-Kaede⁺, lck-GFP⁺, and *rag1⁺* lymphoid populations (Extended Data Fig. 5e-h). Concordantly, *eif3ba*^57^, a key NCC regulator, was detected in zTMT by WISH (Extended Data Fig. 5i). CRISPR/Cas9-mediated knockout^58^ of *eif3ba* significantly reduced fli1a-eGFP⁺ and sox10-GFP⁺ zTMT signals at 5 dpf (Extended Data Fig. 5j-m). These findings jointly revealed that zTMT originates from *sox10^+^* NCCs.

In order to define the process by which *sox10*^+^ NCCs give rise to zTMT, we performed live imaging of *Tg(sox10:mCherry; fli1a:eGFP)* zebrafish embryos. An arc-shaped fli1a-eGFP⁺ structure first emerged adjacent to the caudal pharyngeal arch by 36 hours post-fertilization (hpf) (Fig. 3k). By 2 dpf, these cells had adopted a rounded or oval morphology and were loosely arranged in a cascading configuration (Fig. 3k,l). Between 2.5 and 3 dpf, the cells extended their cellular protrusions to initiate lumen formation (Fig. 3k,l and Supplementary Video 10). The cells became progressively elongated, aligned, and compacted into a semi-coiled, flattened epithelial structure at 3–4 dpf (Fig. 3k,l). After 4 dpf, these cells appeared as a spindle-shaped, tightly packed bundle of fli1a-eGFP⁺ signals that extended ventrally along the BL5 muscle (Fig. 1h,3l). Quantitative analysis revealed that maturation was accompanied by coordinated changes in size and shape. The sagittal length of the zTMT increased from 80.9 μm at 3 dpf to approximately 142 μm by 6 dpf, after which it stabilized (Fig. 3m). Concurrently, the inner diameter expanded from 1.7 μm to 3.6 μm between 3 and 6 dpf (Fig. 3n), while curvature along its ventral segment decreased from 0.030 to 0.016 (Fig. 3o), thus reflecting progressive straightening. Together, these results established that zTMT is derived from *sox10*^+^ NCCs and acquires its mature architecture through a stereotyped morphogenetic program. This program is initiated with loosely organized NCCs, proceeds through lumenogenesis and epithelial compaction, and culminates in a luminalized epithelial conduit that provides the structural foundation for TCPs homing during T lymphopoiesis.

### Sox10 directs morphogenetic remodeling of NCCs in zTMT establishment

We observed that *sox10*^+^ NCCs developed into zTMT, transitioning from a loosely organized, rounded morphology through lumen formation to a tightly packed, spindle-shaped epithelial conduit (Fig. 3k). Sox10 expression persists throughout the morphogenetic process (Fig. 3k). These findings prompted us to investigate the functional requirements of Sox10 in the establishment of zTMT. Sox10 is a core transcription factor required for NCC specification, survival, and differentiation^29^. We generated Sox10 loss-of-function mutants using CRISPR/Cas9-mediated knockout through co-injection of sgRNAs targeting exons 2 and 4 (Extended Data Fig. 6a). A stable mutant line carried a 24-bp deletion in exon 2 and a combined 3-bp deletion plus a 22-bp insertion in exon 4, resulting in reduced *sox10* mRNA levels and a truncated 259-amino acid protein (compared to 485 aa in the wild-type) that was undetectable by quantitative PCR (qPCR) and western blotting (WB) (Fig. 4a,b and Extended Data Fig. 6b). Impressively, *sox10*^⁻/⁻^ larvae displayed a significantly shortened zTMT (mean sagittal length: 116.3 μm in mutant vs. 151.7 μm in siblings) and increased ventral curvature (0.022 vs. 0.017) at 6 dpf (Fig. 4c-e). And the *sox10*^⁻/⁻^ fli1a-eGFP⁺ zTMT cells remained predominantly oval and failed to adopt the elongated, spindle-like morphology observed in control siblings, although the number of fli1a-eGFP⁺ cells for zTMT was strikingly comparable between *sox10*^⁻/⁻^ mutants and their siblings (Fig. 4c,f and Extended Data Fig. 6c). However, mosaic expression of *sox10* under the *fli1a* promoter fully rescued this defect, restoring elongated cell shapes in *sox10*^⁻/⁻^ mutants (Fig. 4g). These results indicated that Sox10 cell-autonomously regulate zTMT cell morphogenesis. Concomitantly, loss of *sox10* compromised zTMT structural integrity, as evidenced by the rapid influx of extraluminally injected dextran into the zTMT lumen and ectopic escape of coro1a-DsRed⁺ TCPs from the lumen (Fig. 4h-m, Extended Data Fig. 6d and Supplementary Videos 11 and 12). Although the migration speed of TCPs was unaffected, thymic cellularity progressively declined (Extended Data Fig. 6e,f). Specifically, the volume of coro1a-DsRed⁺, lck-eGFP^+^, and rag2-DsRed^+^ signals was significantly reduced in *sox10^-/^*^-^mutant thymus at 5 dpf (Fig. 4n,o). These defects were not attributable to altered proliferation (EdU⁺ cell counts) or apoptosis (acridine orange^+^ cell counts) in zTMT cells (Extended Data Fig. 6g,h). Together, these results indicated that Sox10 plays a crucial role in driving the morphogenetic remodeling of zTMT cells from an immature oval shape to an elongated, functionally competent configuration, thereby establishing the structural foundation essential for TCPs homing (Fig. 4p).

**Fig. 4.**
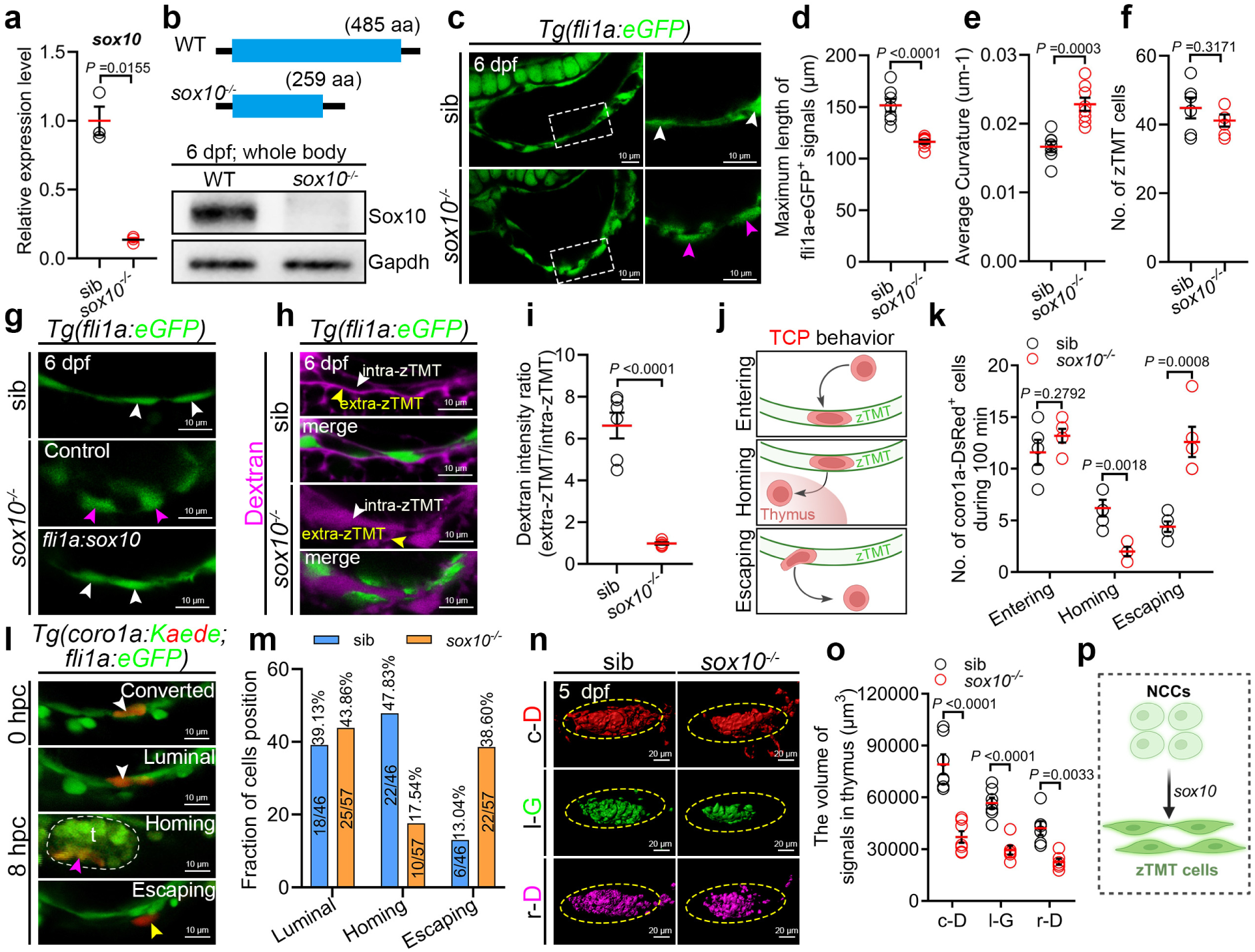
Sox10 regulates the morphogenetic remodeling of zTMT cells. **a**, qPCR analysis of *sox10* transcript levels in sorted *fli1a*^+^*flk1*^-^ cells from the zTMT region of sib and *sox10*^-/-^ zebrafish at 6 dpf. **b**, Schematic diagram (top) and western blot (bottom) showing peptide segments lengths of Sox10 in WT and *sox10*^-/-^ zebrafish. **c**, Confocal images showing morphology of fli1a-eGFP^+^ zTMT in sib and *sox10*^-/-^zebrafish at 6 dpf. Enlarged views of white dashed boxes are shown in the right panels. **d-e**, Quantification of maximum length (**d**) and average curvature (**e**) of the fli1a-eGFP^+^ zTMT in sib and *sox10*^-/-^ zebrafish at 6 dpf. **f**, Quantification of the number of fli1a-eGFP^+^ zTMT structures in sib and *sox10*^-/-^ zebrafish at 6 dpf. **g**, Confocal images showing morphology of fli1a-eGFP^+^ zTMT cells in sib, *sox10*^-/-^, and *sox10^-/-^ /Tg(fli1a:sox10)* zebrafish at 6 dpf. **h**, Confocal images showing localization of Alexa647-dextran^+^ signals in intra-zTMT and extra-zTMT regions in sib and *sox10*^-/-^ zebrafish at 6 dpf. **i**, Quantification of the ratio of extra-zTMT to intra-zTMT dextran intensity in sib and *sox10*^-/-^ zebrafish. **j**, Schematic diagram of TCP behaviors (entering zTMT lumen; homing to the thymus from zTMT lumen; escaping from zTMT lumen to outside) in relation to zTMT. **k**, Quantification of dynamic behaviors of coro1a-DsRed^+^ cells during a 100-min period in sib and *sox10*^-/-^ zebrafish at 5 dpf, according to categories (**j**). **l**, Representative confocal images showing photo-coverted red coro1a-Kaede^+^ cell (white arrowhead) in zTMT lumen at 0 hpc and their position at 8 hpc (white arrowhead, luminal TCP; magenta arrowhead, homing TCP; yellow arrowhead, escaping TCP). **m**, Fraction and number of photo-coverted red coro1a-Kaede^+^ cell in each position category at 8 hpc in sib and *sox10*^-/-^ zebrafish. **n**, Representative 3D images of coro1a-DsRed^+^ (c-D^+^), lck-GFP^+^ (l-G^+^), and rag2-DsRed^+^ (r-D^+^) signals in the thymus (outlined by yellow dashed lines) of sib and *sox10*^-/-^zebrafish at 5 dpf. **o**, Quantification of c-D^+^, l-G^+^, and r-D^+^ signal volumes in the thymus (yellow elliptical dotted lines) of sib and *sox10*^-/-^ zebrafish. **p**, Schematic illustration that NCCs give rise to zTMT through *sox10*-dependent morphogenetic remodeling. WT, wild type; hpc, hour post-conversion. White and magenta arrowheads in **c**,**g** indicate spindle-like and oval morphologies of zTMT cells, respectively. White and magenta arrowheads in (H) indicate intra-zTMT and extra-zTMT regions. Thymus in **l** is outlined by white dashed lines. The experiment in **a** was repeated three times independently. Each dot in **d-f**,**i**,**k**,**o** represents an individual zebrafish larva. Error bars represent mean ± SEM. Unpaired two-tailed Student’s t-test. *P* values are included in the graphs. Schematic diagrams in **j**,**p** were created with BioRender.com.

### Sox10 activates *cdc42* to remodel actin and shape the zTMT

To elucidate the molecular mechanisms by which Sox10 governs zTMT morphogenesis, we performed an in-depth reanalysis of the scRNA-seq data specifically from the zTMT cell population. Given the pivotal role of actin cytoskeleton remodeling in cellular shape dynamics during tissue morphogenesis, we selected Cdc42^35^ as a key candidate. This observation is supported by the significant enrichment of Cdc42-associated pathways in zTMT cells, including Cdc42 protein signal transduction (GO:0032488), small GTPase-mediated signal transduction (GO:0007264), and Rho GTPase binding (GO:0017137) (Fig. 5a). As a Rho family GTPase, Cdc42 orchestrates actin cytoskeletal dynamics to regulate cell polarity, filopodia formation, and directional morphogenesis^35^. qPCR and WB analyses revealed a significant increase in Cdc42 at both the transcript and protein levels from 2 to 4 dpf (Fig. 5b and Extended Data Fig. 7a). Immunofluorescence analysis indicated Cdc42 enrichment specifically within fli1a-eGFP⁺ zTMT cells at 6 dpf (Fig. 5c). Critically, both Cdc42 protein levels and *cdc42* transcript abundance were significantly diminished in the *sox10*^−/−^ mutant zTMT cells (Fig. 5,e and Extended Data Fig. 7b). Bioinformatics analysis (AnimalTFDB4) predicted two conserved Sox10-binding motifs (CTTTGT) in the *cdc42* promoter (Fig. 5f). Chromatin immunoprecipitation (ChIP) using an anti-HA antibody in embryos expressing C-terminally HA-tagged Sox10 manifested specific enrichment of these promoter regions (Fig. 5f). Luciferase reporter assays indicated that Sox10 co-expression significantly enhanced the activity of the wild-type *cdc42* promoter, whereas mutation of its binding sites (notably site 1) abolished this effect (Fig. 5g). Furthermore, mosaic overexpression of *cdc42* under the *fli1a* promoter notably restored the elongated morphology of fli1a-eGFP⁺ zTMT cells in *sox10*^−/−^ larvae, converting their configuration from oval to rod-shaped forms (Fig. 5h and Extended Data Fig. 7c). To test Cdc42 function, we treated embryos with ML141^59^ (a selective, reversible Cdc42 inhibitor) from 3 to 5 dpf (Fig. 5i). ML141 treatment did not significantly affect the embryonic development of wild-type zebrafish (Extended Data Fig. 7d). Notably, this treatment disrupted zTMT barrier integrity, causing dextran to leak from the tissue periphery into the lumen (Fig. 5j and Extended Data Fig. 7e), recapitulating the zTMT cell phenotype of a round morphology in *sox10*^−/−^ mutants (Fig. 5i). Quantitative tracking showed impaired homing of coro1a-DsRed^+^ TCPs to the thymus in ML141-treated larvae (three of eight TCPs, 37.5%) compared with controls (eight of nine TCPs, 88.9%) (Extended Data Fig. 7f,g). The remaining TCPs escaped ectopically from the zTMT (Extended Data Fig. 7f,g and Supplementary Videos 13 and 14). Consequently, the thymic coro1a-DsRed^+^ volume and *rag1* mRNA levels were reduced (Extended Data Fig. 7h). These data established that Cdc42 drives zTMT morphogenesis by enabling actin-dependent cell elongation and establishing a low-permeability lumen barrier.

**Fig. 5.**
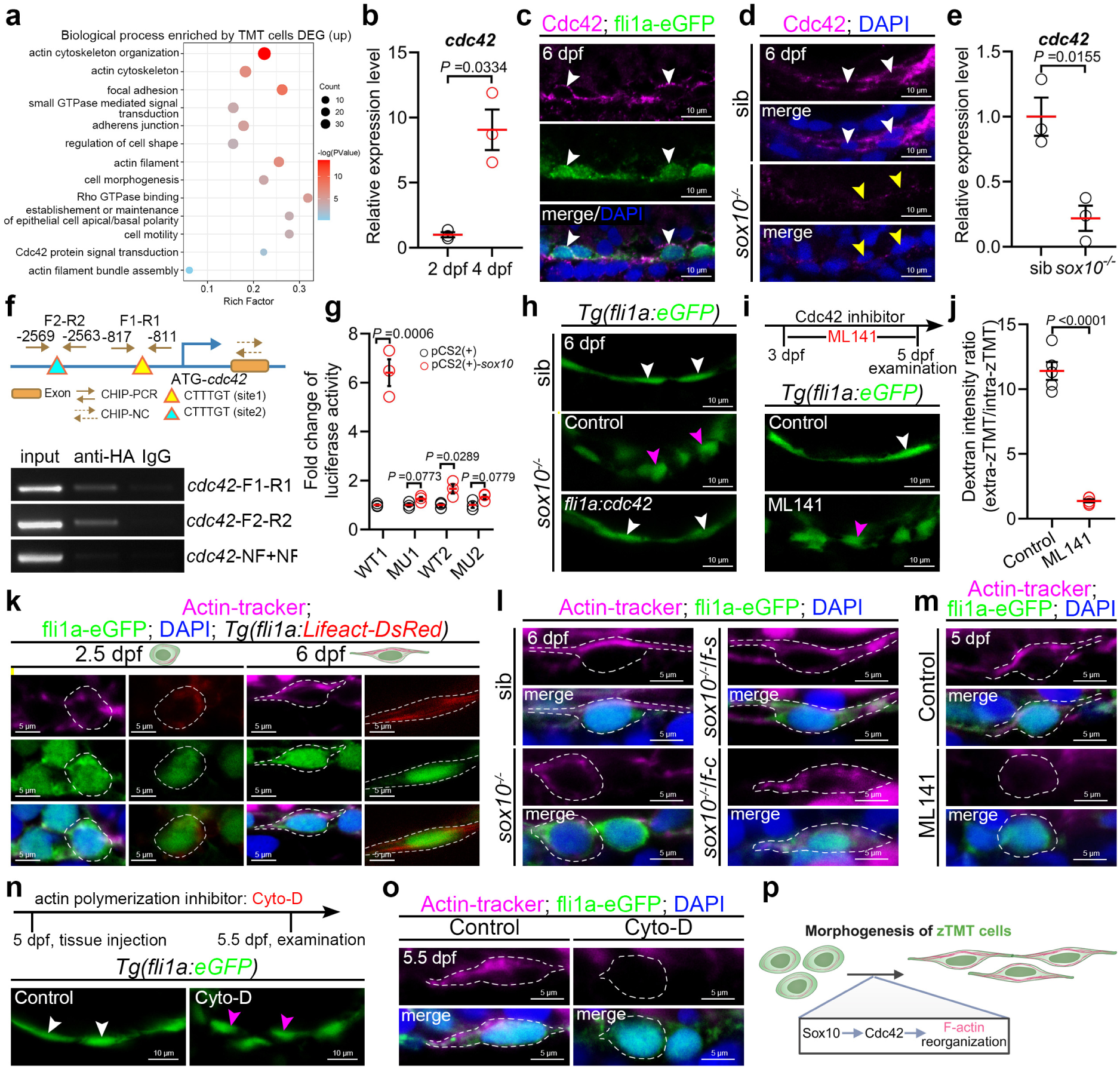
Sox10 activates *cdc42* to remodel actin cytoskeleton and regulate zTMT morphogenesis. **a**, GO analysis of biological process showing enrichment of Cdc42- and cytoskeleton-related terms in cluster 12. **b**, qPCR analysis of *cdc42* transcript levels in sorted *fli1a*^+^*flk1*^-^ cells from the zTMT region of 2 dpf and 4 dpf. **c**, Confocal images showing immunofluorescence staining of Cdc42 co-stained with fli1a-eGFP^+^ zTMT at 6 dpf. **d**, Confocal images showing immunofluorescence of Cdc42 in the zTMT region of sib and *sox10*^-/-^zebrafish at 6 dpf. **e**, qPCR analysis of *cdc42* transcript levels in sib and *sox10*^-/-^ zebrafish at 6 dpf. **f**, Top: Schematic diagram of the predicted *cdc42* promoter with two potential Sox10 binding sites. Bottom: ChIP-PCR gel results. Sites 1 and 2 were amplified using *cdc42*-F1/R1 and *cdc42*-F2/R2 primers, respectively; *cdc42*-NF/NR primers served as negative control. **g**, Quantification of fold changes in the luciferase activities after transfection with indicated vectors. **h**, Confocal images showing morphology of fli1a-eGFP^+^ zTMT in sib, *sox10*^-/-^, and *sox10^-/-^/Tg(fli1a:cdc42)* zebrafish at 6 dpf. **i**, Top: Schematic diagram of the ML141 treatment regimen. Bottom: Confocal images showing morphology of fli1a-eGFP^+^ zTMT in control and ML141-treated zebrafish at 5 dpf. **j**, Quantification of the ratio of extra-zTMT to intra-zTMT dextran intensity in control and ML141-treated zebrafish. **k**, Confocal images showing actin-tracker^+^ (magenta) and fli1a-Lifeact-DsRed^+^ (red) signals in fli1a-eGFP^+^ zTMT cells (green) at 2.5 dpf and 6 dpf. **l**, Confocal images showing actin-tracker^+^ signals (magenta) within fli1a-eGFP^+^ zTMT cells (green) in sib, *sox10*^-/-^, *sox10*^-/-^ /*Tg*(*fli1a*:*sox10)*, and *sox10*^-/-^/*Tg*(*fli1a*:*cdc42)* zebrafish at 6 dpf. **m**, Confocal images showing actin-tracker^+^ signals (magenta) within fli1a-eGFP^+^ zTMT cells (green) in control and ML141-treated zebrafish at 5 dpf. **n**, Top: Schematic diagram of the cytochalasin D (cyto-D) treatment regimen. Bottom: Confocal images showing morphology of fli1a-eGFP^+^ zTMT in control and cyto-D-treated zebrafish at 5.5 dpf. **o**, Confocal images showing actin-tracker^+^ signals (magenta) within fli1a-eGFP^+^ zTMT cells (green) in control and cyto-D-treated zebrafish at 5 dpf. **p**, Schematic illustration of the Sox10-Cdc42-actin axis in zTMT morphogenesis, created with BioRender.com. White arrowheads in **c** indicate the overlapping signals. White and yellow arrowheads in **d** indicate the zTMT cells of sib and *sox10*^-/-^, respectively. White and magenta arrowheads in **h**,**i**,**n** indicate spindle-like and oval morphologies of zTMT cells, respectively. White dashed lines in **k-m**,**o** delineate the zTMT cells. The experiment in **b**,**e**,**g** was repeated three times independently. Each dot in **j** represents an individual zebrafish larva. Error bars represent mean ± SEM. Unpaired two-tailed Student’s t-test. *P* values are included in the graphs.

We extensively investigated how the Sox10-Cdc42 axis orchestrates cytoskeletal remodeling during NCC morphogenesis in the zTMT configuration. scRNA-seq of the zTMT cell population revealed strong enrichment of cytoskeleton-related GO terms, including actin binding (GO:0003779), actin cytoskeleton organization (GO:0030036), actin filament bundle assembly (GO:0051017), cell morphogenesis (GO:0000902), and establishment or maintenance of cell polarity (GO:0007163) (Fig. 5a), implicating actin dynamics as the central driver of zTMT assembly. Direct visualization of F-actin dynamics in *Tg(fli1a:eGFP)* larvae revealed the stereotypical morphogenesis of wild-type zTMT cells. At 2.5 dpf, these cells displayed an oval morphology with weak, diffuse cortical F-actin localization; however, by 6 dpf, they underwent marked elongation, accompanied by a robust reorganization of F-actin into asymmetric bundles oriented along the cellular long axis (Fig. 5k and Extended Data Fig. 7i). *sox10*^−/−^ mutants exhibited severely impaired F-actin polarization through 6 dpf, with F-actin diffusely distributed throughout the cytoplasm, displaying significantly reduced fluorescence intensity, and largely failing to form asymmetric bundles (Fig. 5l). Pharmacological inhibition of Cdc42 with ML141 induced F-actin defects, with the F-actin signal diminishing to near-background levels, diffuse cytoplasmic distribution, and a virtually complete absence of asymmetric bundles (Fig. 5m). Critically, mosaic overexpression of *sox10* and *cdc42* driven by the *fli1a* promoter substantially rescued both F-actin polarization and signal intensity in *sox10*^−/−^ zTMT cells (Fig. 5l). To determine whether F-actin polymerization is required for zTMT morphogenesis, we treated 5 dpf larvae with cytochalasin D^60^, a well-established inhibitor that blocks actin polymerization by capping filament-barbed ends and preventing monomer addition (Fig. 5n). This treatment markedly impaired the assembly of asymmetric F-actin bundles and retained zTMT cells in a rounded morphology persisting through 5.5 dpf, thereby recapitulating both the cytoskeletal abnormalities and the diminished *rag1* mRNA expression in the thymus in the *sox10*^−/−^ mutants and Cdc42-inhibited larvae (Fig. 5n,o and Extended Data Fig. 7j). Collectively, our genetic and pharmacological evidence reveals that Cdc42 is a key effector of Sox10 in orchestrating F-actin reorganization and zTMT cell morphogenesis (Fig. 5p).

### TMT-like structures surround the thymus periphery in embryonic mice

The presence of zTMT in zebrafish prompted us to investigate whether a structurally or functionally similar organ exists in mice. We obtained sagittal sections of tissues surrounding the thymus in embryonic day (E) 13.5–E15.5 mice, when the initial wave of thymic seeding progenitors (TSPs) moves to the thymus^6^. Given the technical challenges of *in vivo* imaging of structures surrounding the murine embryonic thymus, we performed HE staining on paraffin-embedded sagittal and transverse sections. We focused on the thymus and its adjacent tissues in E15.5 mouse embryos because the zTMT extension largely covers the thymus in zebrafish (Extended Data Fig. 2b). Histological examination revealed a well-organized band of flattened cells along the peripheral border of the thymus, arranged in at least a bilaminar configuration and interspersed with several ovoid cells (Fig. 6a,b). Consistently, sagittal SEM revealed flattened cells in the thymic dorsum, where the lymphocyte-like cells were enveloped by flattened cells (Fig. 6c). The end, were stained for CD34 (mesenchymal cells, epithelial, and endothelial cells), CD45 (leukocytes), or IKZF1 (lymphocytes). We observed that CD34^+^ cells with flattened morphology were arranged in a typical tubular pattern along the dorsal edge of the CD45^+^ or IKZF1^+^ thymus (Fig. 6d,e). Within this CD34^+^ structure, several CD45^+^ and IKZF1^+^ cells were detected (Fig. 6d,e). The cellular characteristics of CD34^+^ structures were subsequently investigated by assessing markers specific to vascular (CD31), lymphatic-vascular (LYVE-1), and thymic epithelial (FOXN1) cells. We detected limited colocalization of CD34^+^ structures with CD31^+^, LYVE-1^+^, or FOXN1^+^ signals (Fig. 6f,h). Concordantly, CD34^+^ cells expressed the T-box transcription factor TBX5 (Fig. 6i), analogous to *tbx5a* in zTMT (Extended Data Fig. 5f). These data indicate that flattened CD34^+^ cells form a tubular structure enclosing multiple lymphocytes, similar to the zTMT cells in zebrafish. CD34^+^ cells are neither classical vascular endothelial cells nor thymic epithelial cells, implying that they represent mouse tunnel microtract (mTMT) cells.

**Fig. 6.**
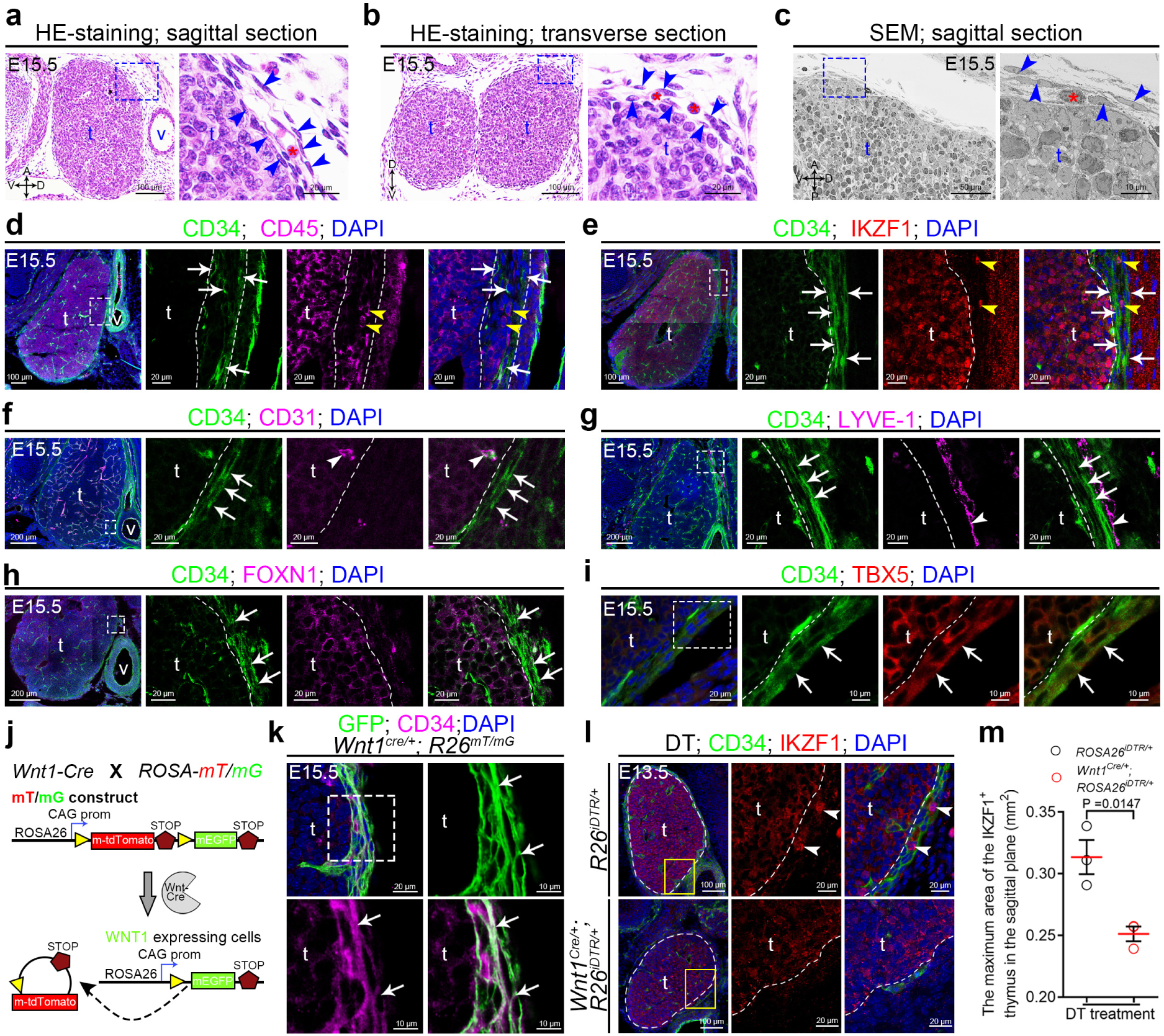
A similar TMT-like structure (mTMT) is present in mouse embryos. **a-c,** Representative HE staining (**a-b**) and SEM images (**c**) of sagittal (**a,c**) and transverse (**b**) sections through the thymus and adjacent tissues in E15.5 mouse embryos. Enlarged views of blue dashed boxes are shown in the right panels. **d-i**, Immunofluorescence images showing the colocalization of CD45 (magenta, **d**), IKZF1 (red, **e**), CD31 (magenta, **f**), LYVE-1 (magenta, **g**), FOXN1 (magenta, **h**), and TBX5 (red, **i**) with CD34 (green) in sagittal sections of E15.5 mouse thymus and adjacent tissues. Enlarged views of white dashed boxes are shown in the right panels. **j,** Schematic picture of the Cre/loxP system used to trace *Wnt1*^+^ NCC-derived lineages. **k,** Immunofluorescence images showing colocalization of CD34 (magenta) with Wnt1-GFP (green) in sagittal sections of E15.5 mouse thymus and adjacent tissues. Enlarged views of white dashed boxes are shown in the right and bottom panels. **l**, Immunofluorescence images of CD34 (green) and IKZF1 (red) in sagittal sections of E13.5 mouse thymus-encompassing tissue after DT treatment. Enlarged views of yellow boxes are shown in the right panels. **m**, Quantification of maximum IKZF1^+^ thymus tissue area (outlined by white dashed lines) following DT treatment. E, embryonic day; v, vein; DT, diphtheria toxin; Blue arrowheads in **a-c** and white arrows in **d-i** mark the path-constituting cells. Red asterisk in **a-c** and yellow arrowheads in **d**,**e** mark cells inside the path. White arrowheads in **f**,**g** indicate CD31^+^ and LYVE-1^+^ signals, respectively. White arrows in **k** indicate the colocalization of GFP^+^ with CD34^+^ signals. White arrowheads in **l** mark the IKZF1^+^cells inside the path. Each dot in **m** represents an individual mouse embryo. Error bars represent mean ± SEM. Unpaired two-tailed Student’s t-test. *P* values are included in the graphs.

### NCCs derived mTMT extends to the thyroid cartilage

Next, we investigated whether the mTMT cells originated from NCCs. We performed genetic lineage tracing using the *Wnt1^Cre^*driver, which labels NCCs from their emergence at E8.5 onward^61^. The *Wnt1^Cre^* transgenic mouse line was used with various indicator transgenic mice to generate recombinants expressing reporter genes in NCC-derived cells. We generated a *Wnt1^Cre^* mouse line using the *ROSA26^mT/mG^*reporter line^62^ (Fig. 6j). GFP fluorescence, indicative of Cre-mediated recombination, was specifically detected in the mTMT regions and colocalized with CD34^+^ signals in the NCCs (Fig. 6k and Extended Data Fig. 8a). CD45^+^ cells were observed within GFP^+^ structures (Extended Data Fig. 8b). To validate the functional requirement of NCC-derived cells for mTMT integrity, we employed a Cre-inducible diphtheria toxin receptor (iDTR) system by breeding *ROSA26^iDTR^* mice with *Wnt1^Cre^*, which enabled DT-induced ablation of *Wnt1*^+^ cells (Extended Data Fig. 8c)^63^. CD34^+^ and SOX10^+^ signals were nearly undetectable in the dorsal boundary of the thymus in DT-treated *Wnt1^Cre^ROSA26^iDTR^* mice compared to DT-treated control *ROSA26^iDTR^* mice, indicating disruption of the mTMT structure in this assay (Extended Data Fig. 8d). Concordantly, the maximum sagittal area of the IKZF1^+^ thymus was 0.25 mm^2^ in DT-treated *Wnt1^Cre^ROSA26^iDTR^*mice, obviously smaller than the 0.311 mm^2^ observed in the *ROSA26^iDTR^* controls (Fig. 6l,m and Extended Data Fig. 8e).

The configuration of mTMT was investigated using this reporter line. HE staining of sagittal sections revealed that the anterior segment of the mTMT extended toward the thyroid cartilage, whereas the posterior segment projected ventrally (Extended Data Fig. 8f). CD34^+^GFP^+^ signals further demonstrated that mTMT encircled the thymus and extended cephalad to the thyroid cartilage (Fig. 7a), suggesting that TCPs enter the thyroid cartilage in mTMT. We were curious about the settlement sites of the TCPs in the mTMT that covered the thymus. Sagittal thymic sections were partitioned into four anatomically defined quadrants relative to the thymic midline: anterior-dorsal (AD), anterior-ventral (AV), posterior-dorsal (PD), and posterior-ventral (PV) (Extended Data Fig. 8g). Quantitative analysis revealed that IKZF1^+^ T cells populated all quadrants, with significantly elevated density specifically in the AD quadrant (Fig. 7b,c). This spatial bias coincided with the anatomical projection of the AD domain toward the thyroid cartilage, suggesting preferential T cell localization in this region (Fig. 7d).

**Fig. 7.**
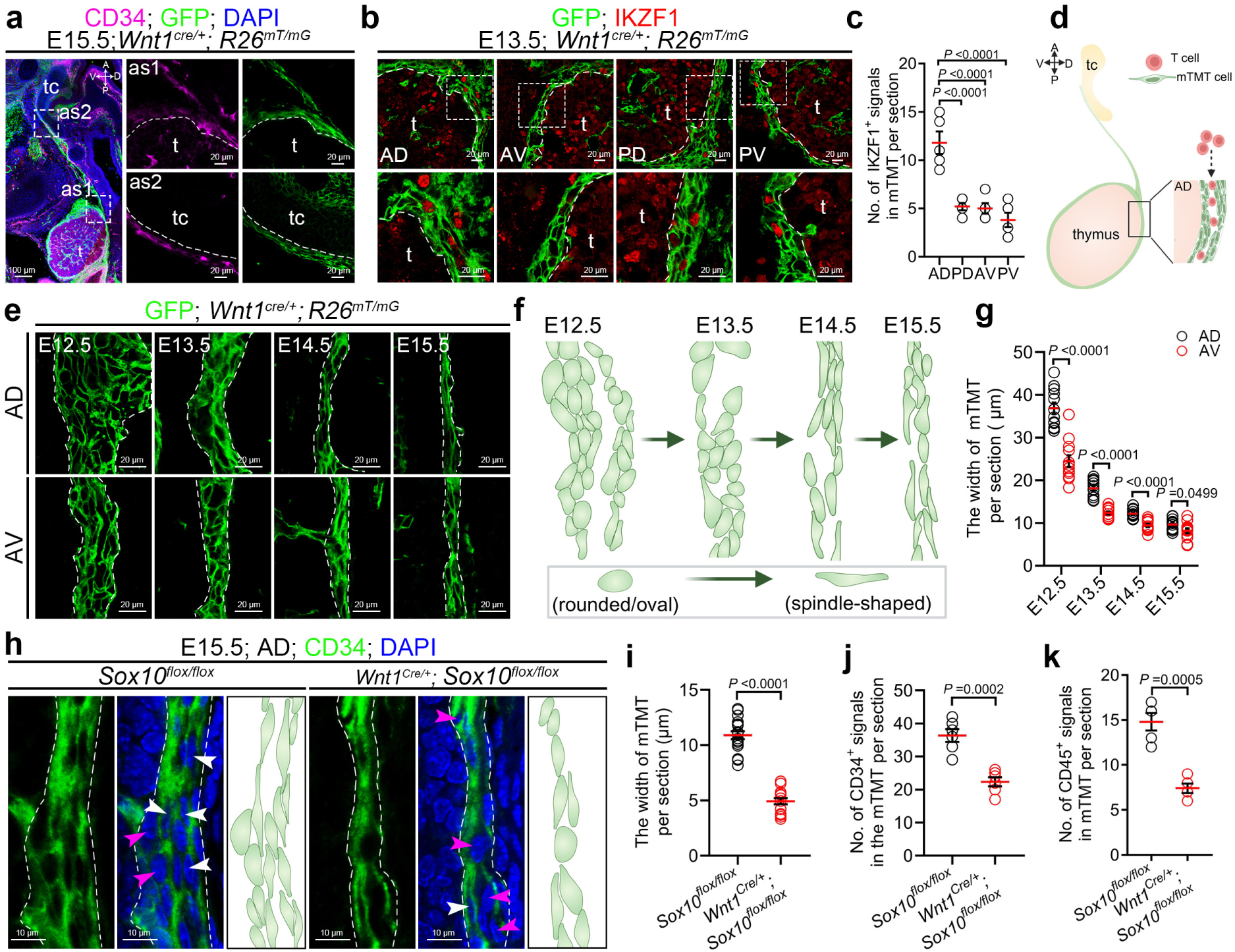
SOX10 regulates morphogenesis of NCC-derived mTMT in mouse embryos. **a**, Immunofluorescence images showing GFP^+^CD34^+^ mTMT encircling the thymus and extending cephalad to the thyroid cartilage. Enlarged views of white dashed boxes are shown in the right panels. **b**. Immunofluorescence images showing localization of IKZF1^+^ signals within GFP^+^ mTMT across the anterior-dorsal (AD), anterior-ventral (AV), posterior-dorsal (PD), and posterior-ventral (PV) regions. Enlarged views of white dashed boxes are shown in the bottom panels. **c**, Quantification of the number of IKZF1^+^ signals in the AD, AV, PD, and PV regions of GFP^+^ mTMT. **d**, Schematic illustration of preferential localization of T cells within mTMT. **e,** Immunofluorescence images showing morphology of GFP^+^ mTMT (outlined by white dashed lines) in AD (top) and AV (bottom) regions at E12.5, E13.5, E14.5, and E15.5. **f**, Schematic illustration of the stereotyped morphogenetic program of mTMT from E12.5 to E15.5. **g**, Quantification of mTMT width in the AD and AV regions across developmental stages. **h**, Immunofluorescence images showing morphology of GFP^+^ mTMT in the AD region of E15.5 *Sox10^flox/flox^* and *Wnt1^Cre/+^; Sox10^flox/flox^* mouse thymus. **i-k**, Quantification of mTMT width (**i**), the number of CD34^+^ signals (**j**), and CD45^+^ signals (**k**) within mTMT in the AD regions of E15.5 *Sox10^flox/flox^*and *Wnt1^Cre/+^; Sox10^flox/flox^* mouse thymus. tc, thyroid cartilage; as, anterior site. White and magenta arrowheads in **h** indicate the spindle-like and oval morphology of mTMT cells, respectively. Each dot in **c**,**g**,**i-k** represents one per section from three mice. Error bars represent mean ± SEM. Unpaired two-tailed Student’s t-test. *P* values are included in the graphs. The schematic diagrams in **d**,**f**,**h** were created with BioRender.com.

### mTMT was produced by NCCs morphogenesis via SOX10-CDC42 axis

The mechanism by which NCCs produce mTMT was investigated. Taking advantage of sequential sections on the reporter lines, we noticed that the GFP^+^ domain progressively narrowed during development, with the dorsal region consistently wider than the ventral region at equivalent stages from E12.5 to E15.5 (Fig. 7e-g and Extended Data Fig. 8g). Concurrently, GFP⁺ cells transitioned morphologically from rounded or oval to spindle-shaped, a pattern that mirrors the developmental trajectory of zTMT, which shifts from rounded/oval configurations at E12.5 to E13.5 to spindle-shaped by E14.5 (Fig. 7e,f). Mechanistically, the conditional deletion of *Sox10* in *Wnt1*-expressing cells (*Wnt1^Cre/+^; Sox10^flox/flox^*) markedly narrowed the CD34^+^ luminal structures within the AD region of the mTMT in sagittal sections relative to *Sox10^flox/flox^* controls (Fig. 7h,I and Extended Data Fig. 8h,i). CD34⁺ cells in mutants predominantly exhibited a subtly ovoid morphology, phenocopying zTMT cells in *sox10^−/−^* zebrafish mutants (Figs. 4g and 7h). Interestingly, whereas *sox10^−/−^* zebrafish mutants displayed unchanged zTMT cell numbers, murine mutants showed a significant reduction in CD34^+^ cell abundance (Figs. 4f and 7j). These cross-species comparisons indicate that *Sox10* governs both cellular morphology and numerical maintenance of mTMT in mice. Concurrently, *Sox10* ablation in NCCs reduced the number of CD45^+^ cells from approximately 15 to 7 per section, reinforcing the functional significance of this path (Fig. 7k and Extended Data Fig. 8j). Immunofluorescence analysis revealed robust Cdc42 protein expression in murine mTMT cells (Extended Data Fig. 8k), which was nearly abolished in *Wnt1^Cre/+^*; *Sox10^flox/flox^* mice (Extended Data Fig. 8l). This genetic evidence establishes Sox10 as a critical upstream regulator of Cdc42 and substantiates the evolutionary conservation of the Sox10–Cdc42 axis during thymocyte migration. Together, these results indicate that the TMT, originating from NCCs, is a conserved organ in both zebrafish and mice and is essential for thymic homing of TCPs during T lymphopoiesis.

## Discussion

Our study has identified a previously uncharacterized organ, the tunnel microtract (TMT), which is a specialized conduit of neural crest origin that enables the homing of T cell progenitors (TCPs) in both zebrafish and mice (Extended Data Fig. 9a). This organ opens our realization of the thymus settlement path for hematopoietic progenitors. Cross-species investigations have revealed anatomical diversity in the TMT. Taking advantage of live images of zebrafish embryos, we noticed that the zTMT is a single-layered tubular structure with a sealed lumen tightly abutting the fifth branchial levator (BL5) muscle. It covers the dorsal surface of the developing thymus anteriorly and terminates posteriorly in a saccular structure close to the kidney medulla (the primary hematopoietic progenitor source, analogous to the mammalian bone marrow)^42^. This anatomical signature favors the movement of TCPs from the kidney to the zTMT via the mesenchyme. However, according to the sectional images, murine mTMT encircles the thymic primordium. Although we extended this path to the thyroid cartilage region and observed that CD45^+^IKZF1^+^ TCPs appeared intensively in the regions adjacent to the thyroid cartilage, we could not reveal the entry process of TCPs into the mTMT from the bone marrow. These issues are challenged by difficulties in live imaging of mice at this stage.

We noticed that zTMT expressed epithelial transcriptional signatures (*epcam*, *krt18a.1*, *krt8*, and *cf1l*) and lacked canonical markers for the vascular endothelium, lymphatic endothelium, or thymic epithelium. Strikingly, zTMT cells express *hoxd11a* and *tbx5a*, factors that govern limb patterning and cardiomyocyte differentiation, suggesting unique signatures of this organ. Concordantly, unlike classical luminal epithelia in the respiratory tract or oviduct, which rely on “9+2” microtubular motile cilia for fluid propulsion^64^, the zTMT consists of a single layer of flattened epithelial cells, non-ciliated cells that form a sealed, sac-like structure with tight junctions and minimal paracellular permeability. Different from canonical conduit epithelia, which function in fluid transport or bulk solute exchange^65^, the unique molecular and structural signature of zTMT appears to support cell migration and development, probably by providing a physically constrained corridor for migrating cells, establishing directional chemotactic gradients via polarized secretion, and mediating transient cell-cell interactions that may deliver cues in T cell development. However, the zTMT lumen exhibited an approximately 8:1 enrichment of lymphoid progenitors (*coro1a^+^*) over myeloid cells (*mpeg1*^+^ macrophages/*lyz*^+^ neutrophils), suggesting that zTMT serves not only as a physical conduit but also as an active cellular sorting compartment. We suspect that the physical properties of the zTMT, with a narrow than ∼5 μm diameter, establish a selective barrier, which permits preferential transit of smaller (3-4 μm), highly deformable TCPs while mechanically restricting larger (5-7 μm), more rigid myeloid cells. However, we could not exclude the possibility that this biophysical filtering mechanism operates synergistically with molecular recognition pathways through a two-tiered selection process wherein the TMT functions as an initial enrichment filter for T lineage-competent cells and the thymic capsule subsequently validates cell fate.

A previous study revealed that zebrafish early progenitors of the aorta-gonad-mesonephros (AGM) extravasate from the vasculature at defined anatomical sites and navigate through the mesenchymal compartment of the pharyngeal arches to reach the thymic epithelial rudiment^18^. Three distinct bloodstream exit sites for migrating progenitors during the embryonic stages are as follows: 1) the triangular vascular network formed by the posterior caudal vein, primary head sinus, and dorsal aorta (the most frequently utilized site); 2) vessels located directly adjacent to the thymic rudiment (the second most prominent exit site); and 3) a rostro-caudal vascular path through the mesenchymal compartment of the pharyngeal arches (the least frequently used route)^18^. Our findings revealed that kidney–derived progenitors migrate to the thymus via the zTMT structure. These TCPs preferentially access the thymus via the caudal end compartment (CEC), a specialized terminal domain of zTMT, rather than through muscle-associated entry sites (MES). A subset of progenitors completely bypasses the zTMT via the ventral exterior route (VER). Our previous studies have suggested heterogeneity in the migratory strategies of TCPs, which may presage divergent lineage commitments in T-cell diversity. Deciphering the navigational logic of these conduits requires the systematic profiling of region-specific guidance molecules, including integrin-ligand interactions^66^, chemokine-receptor pairs^18^, Ephrins^67^, and Semaphorins^67^, to elucidate how spatially patterned cues orchestrate progenitor migration during heterogeneous T cell development.

In addition to its role in trafficking, emerging evidence indicates that zTMT provides an active microenvironment for early T-lineage priming. Single-cell RNA sequencing (scRNA-seq) reveals enrichment of Gene Ontology (GO) terms associated with T cell differentiation (GO:0030217) and hematopoietic progenitor cell differentiation (GO:0002244) (Fig. 3d), while some rag2-DsRed⁺ signals appeared in the lumen of fli1a-eGFP⁺ zTMT structures. Rag2, in concert with Rag1, orchestrates V(D)J recombination, which is essential for T-cell receptor (TCR) diversity, with signalling inputs including the Notch1, IL-7/STAT5, and Wnt/BMP pathways^46, 68, 69^. The probability of the zTMT organ to turn on the expression of Rag2 along TCPs homing suggested the potential TCR rearrangement and β-selection outside the thymus. Similarly, we observed IKZF1^+^ cells in the mouse TMT (mTMT). Therefore, we suspect that the TMT organ initiates the priming of T cells during their movement. This poses a provocative avenue for future investigations to extensively understand extrathymic T cell development in vertebrates.

The neural crest cells (NCCs) of the TMT extend our understanding of NCC contributions to immune development and highlight the crosstalk between the nervous and immune systems. Although NCCs are conventionally recognized as providing thymic stromal support and vascular elements^25^, our findings revealed their additional role in constructing a dedicated TMT conduit through morphogenesis. Extensive studies have established that NCC morphogenesis is a conserved driver of diverse vertebrate organogenesis, encompassing the craniofacial skeleton, enteric nervous system, melanocytes, and adrenal glands^20, 70^. Several pathways are involved in the process. For example, Foxp4-dependent networks are involved in periodontal ligament formation, Neuregulin1-ErbB3 signalling coordinates skeletal muscle progenitor survival and differentiation, NRG1-YAP pathways enabling Schwann cell maturation and myelination, Pth1r-Ihh interactions modulating nasal cartilage hypertrophy and midfacial patterning, and TWIST1-centered chromatin modules (with CHD7, CHD8, and WHSC1) stabilizing early NCC identity, migration, and ectomesenchymal commitment, with relevance to neurocristopathies^71–75^. These recurrent NCC-driven processes of migration, differentiation, and tissue interactions suggest that TMT morphogenesis aligns with established pathways that construct the TMT.

Although NCCs are classically viewed as multipotent mesenchymal progenitors that undergo epithelial-to-mesenchymal transition (EMT) upon delamination to enable migration, accumulating evidence reveals their post-migratory plasticity in adopting epithelial-like states^21^. In trunk lineages, NCCs can re-epithelialize to generate boundary cap cells at the CNS-PNS border or to form perineurial glia with tight junction-like properties^76^. TMT morphogenesis, in which loosely arranged ovoid NCC-derived progenitors progressively adopt spindle-shaped morphologies, align apicobasally, and form tight junctions to create a polarized conduit, recapitulates the key features of mesenchymal-to-epithelial transition (MET)^77^. This reversal of initial EMT supports tissue compaction and lumen formation, as observed during kidney nephron development and somitogenesis^77^.

Mechanistically, Sox10, a core transcriptional regulator of neural crest cell (NCC) fate, is essential for TMT morphogenesis (Extended Data Fig. 9b). In zebrafish *sox10* mutants (*colorless*/*cls*), broad defects in NCC migration and lineage diversification are well documented^78^. However, *fli1a*⁺ zTMT progenitor numbers are largely maintained in *sox10* mutants, while zTMT morphogenesis and epithelial integrity are severely compromised. These results strongly suggest redundant or compensatory pathways that safeguard early NCC specification, survival, and development of the TMT lineage in the absence of Sox10 function. Canonical neural crest signals, including Wnt, Notch, BMP, and FGF, are the leading candidates for such compensation^79^. They likely cooperate with Sox10 to control spatial organization and the critical morphological transition from rounded to spindle-shaped cells that establish a polarized epithelium, a step disrupted in *sox10* mutants.

Notably, Sox10 directly regulates *cdc42* in order to regulate NCC morphogenesis in the zTMT configuration (Extended Data Fig. 9b). Cdc42, a Rho family GTPase, orchestrates F-actin cytoskeletal reorganization that shifts from uniform cytoplasmic distribution at 2–3 dpf to robust apicobasally aligned bundles by 4–6 dpf in zebrafish. This dynamic spatial reorganization is critical for lumen formation and the maintenance of epithelial integrity in developing zTMT. Cdc42 and its downstream F-actin networks have been implicated in the regulation of cell shape and tissue morphogenesis across diverse systems^35^. Localized Cdc42 activation promotes Arp2/3-dependent branched actin polymerization or formin-mediated linear filaments to facilitate apical constriction, filopodia extension, cell elongation, and collective migration during key events such as gastrulation, neural tube closure, and epithelial tubulogenesis in the salivary glands, trachea, kidney, and vascular lumens^37^. Similar Cdc42-dependent mechanisms balance junctional tension via GEFs and GAPs such as ECT2 and ARHGAP12 and coordinate polarity protein positioning through regulators such as KLF5-RICH2 during respiratory tubule branching^80^. These conserved functions align closely with the morphological transitions observed during TMT development in which NCC-derived progenitors shift from ovoid clusters to spindle-shaped cells that support conduit assembly in zebrafish and mice. Our findings extend this paradigm by unveiling Sox10 as an upstream transcriptional regulator of Cdc42 in this NCC-specific context, providing a direct mechanistic link between neural crest fate specification and the cytoskeletal dynamics essential for TMT morphogenesis. Although Cdc42-F-actin signalling is recurrent in endoderm- and mesoderm-derived tubulogenesis in the lungs, kidneys, and salivary glands, its regulation by Sox10 in NCC-derived structures represents a notable adaptation to an immune-related niche.

## Acknowledgments

We thank L. Luo, T. Zhong, Q. Wang for transgenic zebrafish lines, Z. Long, J. Zhang, W. Pan for discussions and technical assistances. We are grateful to Xixia Li and Zhongshuang Lv for helping with sample preparation, taking SEM images at the Center for Biological Imaging (CBI), Institute of Biophysics, Chinese Academy of Science. This work was supported by National Key Research and Development Project (2023YFA1800100), National Natural Science Foundation of China Grants (32470862, 32470869), Chongqing Population and Health Special Funding of China (CSTB2023TIAD-KPX0056), Natural Science Foundation of Chongqing Grants (CSTB2024NSCQ-MSX1222, CSTB2024NSCQ-QCXMX0021), Postdoctoral Fellowship Program of CPSF under Grant Number GZC20232989, and Fundamental Research Funds for the Central Universities (SWU-KQ23014).

## Author contributions

F.Z., J.H. and L.Li developed the concepts and designed the experiments. F.Z. and J.H. performed the experiments and analyzed data. L.X. and Z.L. performed single-cell RNA sequencing and data analysis. L.X. and Y.L. conducted imaging experiments and 3D structural analysis. H.C. and L.D. contributed to tissue section preparation. J.Z. and G.C. performed plasmid construction and microinjection. X.H. assisted with experiments involving hydrogel injection. L. Luo and L. Li provided technical assistance and discussed the results. F.Z. and L. Li wrote the manuscript. All authors read and approved the final manuscript.

## Competing interests

The authors declare no competing interests.

## Methods

### Animals

Zebrafish were bred and maintained under standard laboratory in conditions according with Institutional Animal Care and Use Committee (IACUC) protocols. Embryos were raised in egg water at 28.5 °C and treated with 0.003% 1-phenyl-2-thiourea (PTU; Sigma-Aldrich, P7629) to inhibit pigmentation. The AB wild-type strain was used in this study. Transgenic lines *Tg(coro1a:DsRed)*^81^*, Tg(coro1a:Kaede)*^81^*, Tg(coro1a:DenNTR)*^40^*, Tg(fli1a:eGFP)*^82^*, Tg(rag2:DsRed)*^40^*, Tg(sox10:NTR-mCherry)*^83^*, Tg(flk1:mCherry)*^84^*, Tg(lck:GFP)*^85^*, Tg(prox1:RFP)*^86^*, Tg(cdh17:DsRed)*^87^*, Tg(mpeg1:DsRed)*^88^*, Tg(sox10:GFP)*^89^*, Tg(gata1a:DsRed)*^90^ and *Tg(coro1a:GFP)*^40^ were used. Mice were maintained under specific pathogen-free (SPF) conditions following IACUC guidelines and were socially housed under a 12-h light/dark cycle at 23 °C, with food and water provided ad libitum. The following mice were obtained from the Jackson laboratory: *Wnt1^Cre^* (*H2az2^Tg(Wnt^*^1^*^-cre)11Rth^*Tg(Wnt1-GAL4)11Rth/J, The Jackson Laboratory, 003829)^91^, *ROSA26^mT/mG^*(B6.129(Cg)-*Gt(ROSA)26Sor^tm4(ACTB-tdTomato,-EGFP)Luo/J^*, The Jackson Laboratory, 007676)^92^, *ROSA26^iDTR^*(C57BL/6-*Gt(ROSA)26Sor^tm1(HBEGF)Awai/J^*, The Jackson Laboratory, 007900)^93^, *Sox10^flox/flox^*(CKOCMP-20665-Sox10-B6J-VC, Cyagen Biosciences, S-CKO-17582)^94^. *Wnt1^Cre/+^; ROSA26^mTmG/+^*mice were generated by crossing the *Wnt1^Cre^* with *ROSA26^mT/mG^*. *Wnt1^Cre/+^; ROSA26^iDTR/+^* mice were generated by crossing the *Wnt1^Cre^* with *ROSA26^iDTR^*. Primers used for genomic genotyping are provided in Supplementary Table 1. All animal experiments were approved by the Institutional Review Board of Southwest University and Chongqing Institute of Green and Intelligent Technology, Chinese Academy of Science (cigit2024003, Chongqing, China).

### Generation of transgenic zebrafish lines

The *Tg(sox10:Cre)*, *Tg(fli1a:loxP-mCherry-loxP-eGFP), Tg(snorc:GFP)*, *Tg(fli1a:Lifeact-DsRed)* and *Tg(tbx5a:nsfb-mCherry)* transgenic zebrafish lines were generated in this study. Transgenic constructs were generated using the Tol2 transposon system^81^. To generate the *pTol2-sox10-Cre* construct, the *sox10* promoter was amplified from the *pDEST-sox10-mCherry* plasmid and subcloned into the *pTol2-cryaa-mCherry* vector using EcoRI and SalI restriction sites, yielding a *pTol2-sox10-cryaa-mCherry* plasmid. Subsequently, the Cre recombinase coding sequence was amplified from the *pTol2-hsp70l-Cre-cryaa-Cerulean* backbone and fused in-frame downstream of the *sox10* promoter. To construct of the *pTol2-fli1a-loxP-mCherry-loxP-GFP* plasmid, the *loxP-mCherry-loxP-eGFP* cassette was amplified from the *pTol2-hsp70l-loxP-mCherry-STOP-loxP-H2B-GFP-cryaa-Cerulean* plasmid and subcloned into a *pTol2-fli1a* vector. For generating the *Tg(fli1a:Lifeact-DsRed)* plasmid, the *Lifeact-DsRed* cassette was amplified from the *pCRII-flk1-Lifeact-DsRed* plasmid^95^ and subcloned into a *pTol2-fli1a* vector. To construct of the *Tg(snorc:GFP)* and *Tg(tbx5a:nsfb-mCherry)* plasmid, cassette of 6.3 and 5.9 kb upstream of the start codon of *snorc* and *tbx5a* genes was respectively amplified as their promoter, following subcloned into *pTol2-GFP* and *pTol2-nfsB-P2A-mCherry* vectors. The resulting plasmids were co-injected with transposase mRNA into one-cell-stage embryos of the AB wild-type zebrafish strain. Embryos exhibiting specific mCherry expression in tissues including the eyes and vasculature were selected and raised to adulthood to establish heritable transgenic lines.

### Generation of *hoxd11a* knock-in reporter line

To generate a *hoxd11a* reporter line (*Ki(Hoxd11a-P2A-Gal4-Vp64;uas:nfsB-P2A-mCherry)*) we used a CRISPSR/Cas9-mediated knock-in strategy. The *hoxd11a* sgRNA, targeted sites within the first exon, was injected at a concentration of 50 ng/μl together with the UgRNA (50 ng/μl), Cas9 mRNA (500 ng/μl) and the donor vector (50 ng/μl; constructed by the China Zebrafish Resource Center, National Aquatic Biological Resource Center, CZRC/NABRC). The founder embryos were raised, and next-generation (F1) progenies with mCherry expression were identified by genotyping for homologous recombination analysis.

### CRISPR/Cas9-mediated knockout

Efficient knockout of the *eif3ba* gene in the F0 generation was achieved by co-injection of three distinct single-guide RNAs (sgRNAs), as previously described^58^. Briefly, three CRISPR RNAs (crRNAs) targeting distinct loci within the *eif3ba* genomic sequence (crRNA-1: AGAGCCTTCGTTCAGTGATC; crRNA-2: CAAGGAGCCTATTGAAGTGG; crRNA-3: ACCAAGTCACGCTCATGAAG) were designed using the predesigned crRNA database from Integrated DNA Technologies (IDT). Each crRNA was annealed with trans-activating CRISPR RNA (tracrRNA) in Duplex Buffer to generate individual sgRNAs. Subsequently, each sgRNA was separately combined with Cas9 protein (Integrated DNA Technologies, 1081059) to generate three distinct ribonucleoprotein (RNP) complexes. These RNP complexes were mixed at an equimolar ratio (1:1:1) to target the *eif3ba* gene. One-cell-stage wild-type AB embryos were injected with the pooled RNP complexes to induce *eif3ba* mutations in the F0 generation. Knockout efficiency was assessed by quantitative PCR (qPCR) analysis of *eif3ba* transcript level using the primers 5’-CGGAGAACATGGAGGCCGAA-3’ and 5’-CAGGGTAGAACTCATTGGTG-3’. The reported phenotype of *eif3ba* mutant zebrafish^57^, characterized by subtle hypopigmentation and altered head morphology, was used to evaluate gene-editing efficiency.

### Photoconversion, and *in vivo* time-lapse imaging

To investigate the contribution of progenitor cells from the caudal hematopoietic tissue (CHT), aorta-gonad-mesonephros (AGM) region, and kidney to thymic T cells, coro1a-Kaede^+^ cells in these tissues of 5 dpf *Tg(coro1a:Kaede)* zebrafish were selected using the ROI mode. Selected cells were irradiated with a 405-nm laser at 50% power for 30 s to induce the photoconversion of green Kaede protein to red color. The number of red coro1a-Kaede^+^ cells in the thymus was quantified at 5 hours post-conversion (hpc). To examine the dynamics of cellular entry and exit within the zebrafish tunnel microtract (zTMT), coro1a-Kaede^+^ cells in the kidney of 5 dpf *Tg(coro1a:Kaede)* embryos were subjected to a 405 nm ultraviolet (UV) laser of 30 s. Subsequently, *in vivo* time-lapse imaging was performed.

*In vivo* time-lapse imaging was executed as previously described^81^. Briefly, the anesthetized embryos were embedded in 1.5% low-melting-agarose and oriented laterally in a 15-mm dish to expose the thymic region. Time-lapse sequences were acquired using a Zeiss LSM700 confocal microscope equipped with a 20× objective. *z*-stacks were acquired with a step size of 1.5 μm, which were recorded at 2-to 3-min intervals over several hours.

### Microinjection of Dextran and PEG hydrogel

For dextran microinjection, Alexa Fluor 647-conjugated dextran (10 kDa; Thermo Fisher Scientific, D22914) was directly injected into tissues adjacent to the caudal musculature and the ventral region of the otic vesicle. Microinjections were performed using a glass capillary needle under a stereomicroscope (Stemi 2000-C, ZEISS). Polyethylene glycol (PEG) hydrogels were formed via covalent cross-linking between amine and *N*-hydroxysuccinimide (NHS) ester groups^44^. Tetra-arm PEG succinimidyl glutarate (Tetra-PEG-SG, *M*_w_ = 10 kDa) and Tetra-arm PEG amine (Terra-PEG-NH_2_, *M*_w_ = 10 kDa) were purchased from SINOPEG (Fujian, China). Cyanine 5 amine (Cy5-NH_2_) was obtained from ApexBio Technology (Beijing, China). To synthesize Cy5-labelled hydrogels, two solutions were prepared separately. Solution A was prepared by dissolving Cy5-NH_2_ (653.77 g/mol) in PBS (pH 7.4) to a final concentration of 2.68 μg/μl. Subsequently, 2.5 μl of this solution was mixed with 1 mg of Tetra-PEG-SG and allowed to react in the dark at room temperature for 10 min. Solution B was prepared by dissolving 0.8 mg of Tetra-PEG-NH_2_ dissolved in 2.5 μl PBS (pH 7.4) and mixing thoroughly at room temperature. To induce hydrogel occlusion *in vivo*, equal volumes of solution A and solution B were sequentially injected into the same target sites in zebrafish. Gelation occurred within 1-2 s upon mixing of the two solutions.

### Chemical treatment

Metronidazole (MTZ) treatment was administered as previously described^40^. *Tg(sox10:NTR-mCherry)* zebrafish were treated with either 0.2% DMSO or 20 mM MTZ (Sigma-Aldrich, PHR1052) dissolved in 0.2% DMSO from 3 to 5 dpf to induce neural crest cells (NCCs) ablation. *Tg(coro1a:DenNTR)* zebrafish were treated with either 0.2% DMSO or 2 mM MTZ (dissolved in 0.2% DMSO) from 5.5 to 8 dpf to induce thymic T lymphocyte ablation. Following treatment, zebrafish were extensively washed and allowed to recover in PTU solution.

ML141^59^(Selleck, S7686) powder was dissolved in DMSO to prepare a 50 mM stock solution. Larvae were incubated in 7.5 µM ML141 working solution from 3 dpf to 5 dpf. Control larvae were incubated in egg water containing 0.15% DMSO and PTU.

Cytochalasin D^60^ (Yeasen, 22144-77-0) powder was dissolved in DMSO to prepare a 2.5mM stock solution. 1 mM Cytochalasin D working solution was directly injected into tissues adjacent to the caudal musculature and the ventral region of the otic vesicle at 5 dpf. Control larvae were injected with PBS.

Diphtheria toxin (DT; Sigma, D0564) treatment was performed as previously reported^63^. To ablate NCCs in mice, *Wnt1^Cre^* mice were crossed with *ROSA26^iDTR^*mice. Pregnant females were administered with diphtheria toxin (20 mg/kg body weight in sterile saline) via intraperitoneal injection beginning at embryonic day 9.5 (E9.5) and continuing once daily for 4 days. Fetuses were collected at E13.5 for genotyping. Identified *ROSA26^iDTR^* and *Wnt1^Cre^ROSA26^iDTR^*embryos were used as control and experimental groups, respectively.

### Whole-mount *in situ* hybridization (WISH), fluorescence *in situ* hybridization (FISH) and immunofluorescence staining

WISH and FISH were performed according to standard protocols^81^. The RNA probes (*ikzf1*, *foxn1*, *rag1*, *fli1a*, *clic2*, *ucmab*, *pah*, *otos*, *stmn1a*, *tbx5a*, *hoxd11a*, *sox9a*, *sox10*, *eif3ba*, *epcam*, *krt18a.1*, *lgals2a*, *dlx4a*, *msx1b*, *sostdc1a*, *cyp1c2*, *lhx9*, *tfr1a*, and *sfrp2*) were transcribed *in vitro* using T3 or T7 RNA polymerase (Thermo Fisher Scientific, EP0101/ EP0111) with Digoxigenin RNA Labelling (Roche, 11277073910). Primers used for probe synthesis are listed in Supplementary Table 1. The Leica VT1000S and Leica CM1950 instruments were used to prepare vibratome sections of zebrafish tissues and frozen sections of mouse tissues, respectively, as previously described^81^. For immunofluorescence staining, tissue pieces were rinsed in PBDT (PBS containing 1% BSA, 1% DMSO, 0.5% TrintonX-100) and blocked with buffer (2% FBS and 0.1% Tween20 in PBS) at room temperature for 1 hour. The primary antibodies were utilized: goat anti-GFP (1:400, Abcam, ab6658), rabbit anti-DsRed (1:400, Takara, 632496), Mouse anti-II-II6B3 (1:400, DSHB, AB_528165); Mouse anti-S58 (1:400, DSHB, AB_528377); Mouse anti-F310 (1:400, DSHB, AB_531863); rabbit anti-SOX10 (1:400, Abcam, ab155279), rabbit anti-FOXN1 (1:400, UABIO, ER1908-98), goat anti-LYVE-1 (1:400, R&D Systems, AF2125), mouse anti-CD31 (1:400, HUABIO, M1511-8), rat anti-CD34 (1:400, BD Biosciences, 553731); mouse anti-CD45 (1:400, proteintech, 60287-1-Ig); rabbit anti-CD45 (1:400, Cell Signaling Technology, 70257); rabbit anti-SOX10 (1:400, Abcam, AB_2650603); rabbit anti-CDC42 (1:400, Huabio, AB_3070234); rabbit anti-SOX9 (1:400, Abcam, AB_2728660); rabbit anti-IKZF1 (1:400, Huabio, AB_2124704); Lcp1 antibody (1:400, GeneTex, AB_11167454); rabbit anti-TBX5 (1:400, Novus, AB_11018767); rabbit anti-HOXD11 (1:400, Novus, AB_3343027). The samples were subsequently washed with PBDT and stained with AF488/555/647-conjugated secondary antibodies: 488 goat anti-Rat (1:400, Abcam, ab150165), 488 donkey anti-Goat (1:400, Thermo Fisher Scientific, A-11055), 555 donkey anti-Rabbit (1:400, Thermo Fisher Scientific, A-31572), 647 donkey anti-Mouse (1:400, Thermo Fisher Scientific, A-31571) at 37°C for 3 hours. For actin staining, tissue samples were incubated with Actin-Tracker Deep Red^96^ (Beyotime, C2209S) for 1 hour following secondary antibodies treated. The WISH signals were captured using a Carl Zeiss Discovery.V20 microscope (Carl Zeiss) and the fluorescence signals were acquired using an LSM880 confocal microscope (Carl Zeiss).

### Western blotting (WB)

The antibodies utilized in this study: rabbit anti-CDC42 (Huabio, AB_3070234); mouse anti-GAPDH (Proteintech, AB_2107436); rabbit anti-HOXD11 (Novus, AB_3343027). The anti-HOXD11 were utilized to detect the protein level of Hoxd11 in wild-type (WT) and *Tg(hoxd11a:nfsB-mCherry)* zebrafish at 3 dpf. Anti-CDC42 were used to evaluate the protein level of actin in WT and *sox10^-/-^* zebrafish at 6 dpf. Detections were completed by Goat anti-mouse or anti-rabbit HRP-conjugated secondary antibodies (Thermo Fisher Scientific, AB_228307/ AB_1965959), followed by enhanced chemiluminescence (ECL Plus). Each protein sample was prepared from approximately 50 zebrafish embryos. All WB experiments were performed independently at least three times with similar results.

### Electron microscopy

Zebrafish samples from the targeted regions (tissue from the region of zebrafish larvae between the posterior portion of the eye and the anterior portion of the yolk sac) were isolated. To perform transmission electron microscope (TEM), the samples were fixed in in a solution containing 2.5% glutaraldehyde, and 2% formaldehyde in Cacodylate buffer (0.1 M cacodylate in water, pH7.0) at 4 ℃ overnight. Following a rinse in the cacodylate buffer, the tissue was dehydrated using a graded series of ethanol to propylene oxide. The tissue was subsequently infiltrated and embedded in epoxy resin and polymerized at 70 ℃ overnight. Semi-thin sections (0.5 mm thickness) were stained with toluidine blue and examined via light microscope. The ultrathin sections (80 nm) were prepared and examined using a transmission electron microscope (Hitachi H-600 TEM) to acquire the TEM images, facilitated by the Sci-Tech Innovation Center at Chongqing Medical University. To perform the scanning electron microscopy (SEM) assay, the samples were fixed with 2.5% glutaraldehyde and 2% paraformaldehyde with PBS. After washing several times, tissues were incubated in the filtered 1% thiocarbohydrazide (TCH) aqueous solution (Sigma-Aldrich, 223220) at room temperature for 30 min, 1% unbuffered OsO4 aqueous solution at 4℃ for 1h and 1% UA aqueous solution at room temperature for 2 h sequentially. Then the tissues were dehydrated through the graded alcohol (30, 50, 70, 80, 90, 100%, 100%, 10min each, at 4℃) into pure acetone. These samples were infiltrated with graded mixtures of acetone and SPI-PON812 resin (composition:21 ml SPI-PON812 (SPI, INC., 02659-AB), 13 ml DDSA (Dodecenyl Succinic Anhydride, SPI, INC., 02827-AF), 11ml NMA (Nadic Methyl Anhydride, SPI, INC., 02828-AF and 1.5% BDMA (N, N-Dimethylbenzylamine, SPI, INC., 02821) using sequential ratios of 3:1, 1:1, and 1:3, followed by pure resin immersion. These tissues were then embedded in pure resin and polymerized for 12 h at 45°C, 48 h at 60°C. The ultrathin sections (70 nm thickness) were sectioned using a ultramicrotome (Leica EM UC6, Germany) and examined by a scanning electron microscope (FEI Helios Nanolab 600i dual-beam SEM) with its an immersion high magnification mode (concentric backscattered (CBS) detector, 2kV, 0.69nA) to acquire the SEM images (backscattered electron signal) by an automated imaging software (AutoSEE).

### Hematoxylin and eosin (HE) staining

Zebrafish larvae and mouse embryos were fixed overnight in 4% paraformaldehyde (PFA) at 4°C. Samples were subsequently dehydrated through a graded ethanol series (70%, 80%, 90%, 95%, and 100%), cleared in xylene, and embedded in paraffin wax. Paraffin blocks were sectioned at a thickness of 10 μm using a Leica RM2016 microtome. Sections were mounted onto glass slides, deparaffinized, rehydrated through a descending ethanol series, and stained with hematoxylin and eosin according to standard protocols^97^. After staining, sections were dehydrated, cleared, and mounted with a permanent mounting medium. Images were captured under a bright-field microscope.

### Three-dimensional (3D) reconstruction and quantification of thymic T cell volume

3D reconstruction was performed using Imaris software (Bitplane) to visualize the fli1a-eGFP^+^ TMT structure and to track the migratory trajectories of coro1a-DsRed^+^ and red coro1a-Kaede^+^ cells into the caudal end compartment (CEC) region of zTMT. For cell size quantification, *z*-stack confocal images of coro1a-DsRed^+^, coro1a-Kaede^+^, rag2-DsRed^+^ and lck-GFP^+^ cells in the thymic regions were processed to generate 3D surface renderings. The total volume of these fluorescently labelled T cell populations in the thymus was measured using the Surface module in Imaris and served as a quantitative readout of thymic T cell development.

### Quantitative measurements of zTMT length and depth

Both transverse and sagittal sections were prepared to assess the anatomical dimensions of the TMT. For depth measurements, transverse sections were prepared at four anatomical landmarks: site 1 (rostral edge of the thymus), site 2 (mid-thymic region), site 3 (the central portion of the BL5 muscle), and site 4 (caudal terminus of the BL5 muscle). Specifically, measurements were taken from the pharyngeal arch to the external region of the thymus (attaching to the dorsal thymic region at sites 1 and 2), from the pharyngeal arch to the external region of the BL5 muscle (attaching to the ventral thymic region at site 3), and surrounding the entire ventral margin of the BL5 muscle at site 4, as defined by the extension of the fli1a-eGFP⁺ signal. For length measurement, sagittal sections were acquired at defined *z*-axis depths (15 μm, 30 μm, 45 μm, and 60 μm). In each plane, the sagittal length was quantified by measuring the distance of continuous fli1a-eGFP^+^ signals extending from the dorso-rostral side of the thymus (*z* = 15 μm, 30 μm, 45 μm) or the region adjacent to the pharyngeal arch (*z* = 60 μm) to the caudal-ventral border of the BL5 muscle. At each site, the depth of the zTMT was defined by the fli1a-eGFP⁺ signal. All measurements were performed on high-resolution confocal images and quantified using Imaris or ImageJ.

### ChIP and luciferase reporter assay

The chromatin immunoprecipitation (ChIP) assay was performed as previously described^81^. Briefly, the *pCS2-HA-sox10* were initially generated and microinjected into WT zebrafish embryos at one-cell-stage. The DNA was extracted at 3 dpf by Chromatin Immunoprecipitation Kit (Sigma-Aldrich, 17-295) and then precipitated by immunoglobulin G (lgG as controls; Thermo Fisher Scientific, 31460) coupled beads (Thermo Fisher Scientific, 10002D) and HA Magnetic Beads antibodies (Beyotime, P2121). PCR was performed by using the designed primers that targeted the predicted Sox10 binding sites. Luciferase reporter constructs (PGL3) containing WT (site1/2: CTTTGT) and mutant (deletion) binding sites within *cdc42* promoter regions were generated. Then, the *PGL3-cdc42* (WT, 200 ng) or *PGL3*-*cdc42* (MU1/2, 200 ng) plasmids, together with *pCS2-sox10* (200 ng) and pRL-CMV (100 ng) plasmid, were transfected into 293T cell lines via Lipo8000(Beyotime, C0533-1.5). PGL3-basic plasmid was used as a control. After 2 days of culture, cells were lysed, and the supernatant was collected by centrifugation at 12,000 × g. Fluorescence intensity was measured in the collected supernatant.

### Bulk RNA Sequencing and data analysis

Suspensions of coro1a-DsRed⁺ cells were collected from the thymus, intra-zTMT, and extra-zTMT regions of zebrafish larvae according to previously described protocols^81^. Approximately 500 coro1a-DsRed⁺ cells from each region were sorted by flow cytometry (Moflo XDP, Beckman) for cDNA library preparation by Anoroad Corporation. RNA sequencing was performed using a PE100 strategy (HiSeq 2500 Illumina) and sequencing reads were aligned to the zebrafish reference genome (Danio rerio, GRCz11). Genes with an adjusted *P* value < 0.1 and log2 fold change > 0 were defined as differentially expressed genes (DEGs) using DESeq2. Focused genes expression of each group was visualized using Heatmap. Gene ontology (GO) biological process enrichment analysis of DEGs was performed using DAVID Bioinformatics Resources (v6.8).

### Single-cell RNA sequencing (scRNA-seq) library preparation and analysis

Fluorescence-activated cell sorting (FACS) was used to isolate fli1a-eGFP^+^flk1-mCherry^-^ cells, which were subsequently loaded into the channels of a Chromium Single Cell G Chip (v3.1 chemistry, PN-1000120). Single-cell libraries were prepared using the Chromium Single Cell 3’ Reagent Kits (v3.1 chemistry PN-1000121) by curaMed Medical Technology Co., LTD (Shanghai, China). Sequencing data were aligned to the zebrafish reference genome (Danio rerio, GRCz11) using Cell Ranger 5.0.0. The resulting matrix was processed with Seurat v3 for the quality control, filtering, dimensionality reduction, clustering analysis, and DEGs analysis. Cluster-specific marker genes and focused genes were visualized using Heatmap and FeaturePlot. GO biological process enrichment analysis was performed using DAVID Bioinformatics Resources (v6.8).

### Developmental Pseudotime Analysis

Single-cell pseudotime trajectory analysis was performed using the R package Monocle3 (v1.2.9) to infer developmental transitions of neural crest cells (NCCs) cluster. Clustering and dimensionality reduction results generated in Seurat were used as input for Monocle3 analysis. The learn_graph function was applied to fit a principal graph within each partition and to reconstruct developmental trajectories with multiple branches. The root node of the trajectory was manually specified based on biological prior knowledge. Cells along the trajectory were ordered in pseudotime and visualized using the plot_cells function. Gene expression dynamics along pseudotime were displayed as heatmaps, in which genes of interest were ordered according to pseudotime and visualized using the heatmap package.

### Image acquisition and processing

Immunofluorescence-stained samples and live zebrafish embryos were imaged using a Zeiss LSM880 or LSM700 confocal microscope. WISH and HE (Sangon biotech, E607318-0200)–stained samples were imaged with a Nikon Eclipse Ni microscope equipped with NIS-Elements BR software (version 5.01). Image annotations, including figure labels, text, arrows, and outlines, were generated using Adobe Photoshop CC 2020. The schematic diagrams and flowcharts presented in the figure were generated using BioRender.com or Adobe Photoshop CC 2020. Time-lapse imaging data were processed using Imaris or ImageJ and subsequently converted into movies with Adobe Premiere Pro CC 2020.

### Statistical analysis

All statistical analyses were performed using two-tailed Student’s *t*-tests in GraphPad Prism (version 9.0). Centre values represent the mean, and error bars indicate the standard error of the mean (S.E.M.). Details regarding sample sizes, error bars, and exact *P* values were provided in the figures and figure legends. All experiments were independently repeated at least three times, and representative images from these replicates are shown.

## Data availability

Raw and source data were provided with this paper. The raw bulk RNA-seq and single-cell RNA-seq (scRNA-seq) data have been deposited in the Gene Expression Omnibus (GEO) under accession numbers GSE319391 and GSE297957, respectively. Additional data supporting the findings of this study, including zebrafish lines, plasmids generated in this work, and chemical reagents, were available within the article and its Supplementary Information, or from the corresponding author upon reasonable request.

## Notes

### Competing Interest Statement

The authors have declared no competing interest.

